# CRISPR-enhanced assessment of variants of unknown significance nominates oncology therapeutic targets and drug repositioning opportunities

**DOI:** 10.64898/2026.01.20.700565

**Authors:** Aurora Savino, Athanasios Oikonomou, Francesca Perrone, Riccardo R De Lucia, Letizia De Pietri, Yousra Belattar, Luke Brown, Miguel L. Grau, Katrina McCarten, Hanna Najgebauer, Umberto Perron, Luca Azzolin, Alexandra Livanova, Paolo Cremaschi, Nuria Lopez-Bigas, Andrea Sottoriva, Matthew A. Coelho, Francesco Iorio

**Affiliations:** Computational Biology Research Centre, Human Technopole, Milan, Italy; Wellcome Sanger Institute, Wellcome Genome Campus, Hinxton, Cambridge, UK; Institute for Research in Biomedicine (IRB Barcelona), The Barcelona Institute of Science and Technology, Barcelona, 08028, Spain; Open Targets, Wellcome Genome Campus, Hinxton, Cambridge, UK; European Molecular Biology Laboratory, European Bioinformatics Institute, Wellcome Genome Campus, Cambridge, UK; Centro de Investigación Biomédica en Red en Cáncer (CIBERONC), Instituto de Salud Carlos III, Madrid, 28029, Spain; Department of Medicine and Life Sciences, Universitat Pompeu Fabra, Barcelona, 08003, Spain; Institució Catalana de Recerca i Estudis Avançats (ICREA), Barcelona, 08010, Spain

## Abstract

Interpreting infrequent somatic variants remains a challenge in cancer genomics. We developed CRISPR-VUS, a framework that uses public Cancer Dependency Map data to identify Dependency-Associated Mutations (DAMs) - variants linked to increased host-gene dependency - with resolution extending to singleton events. Analysis of 977 cell lines across 36 cancer types identified 2,376 DAMs in 1,383 genes, including 1,260 not established as cancer drivers. DAM-bearing genes converge on oncogenic networks, while recurrence in histology-matched tumours, functional-impact predictions, tractability and pharmacological associations enable systematic prioritisation. Prime editing showed that the prioritised NSCLC-specific RTN4IP1-A80T DAM conferred a significant competitive growth advantage in a lung epithelial model, nominating a candidate driver allele. Exploratory pharmacological testing showed a greater maximal response to istaroxime in ATP1B3-I189M-bearing than in ATP1B3-wild-type cells. CRISPR-VUS combines dependency-based rare-variant discovery with evidence-guided prioritisation to nominate candidate drivers, therapeutic targets and drug-repositioning hypotheses. Interactive results are available at https://vus-portal.fht.org/.

## Introduction

Cancer is driven by somatic mutations that confer selective advantages and enable malignant phenotypes^1,2^.

Identifying cancer driver genes is a central goal of cancer genomics and has been pivotal in the development of targeted therapies^3–5^. Many of these therapies exploit oncogene addiction, whereby cancer cells harbouring a tumour-promoting alteration in a key oncogene become vitally dependent on its activity^6–10^.

Conventional driver-discovery approaches exploit recurrence across patient cohorts^11^, supplemented by evolutionary conservation, predicted functional impact and pathway information^12,13^. More recently, machine-learning and ensemble-based methods, such as CancerVar^14^, REVEL^15^ and CADD^16^, have improved variant interpretation by integrating sequence, functional and clinical annotations. Nevertheless, these methods infer relevance from existing evidence or features predictive of general deleteriousness. Their evidence bases and training or reference datasets also provide substantially more information for previously characterised variants and genes. Consequently, they may leave rare, unannotated or lineage-specific somatic variants unresolved, particularly when their effects depend on cellular context^17,18^.

Cancer cells can become dependent not only on activated oncogenes but also on genes required to tolerate oncogenic or cellular stress, generating oncogene and non-oncogene addictions^19^. Large-scale RNAi and CRISPR-Cas9 viability screens have mapped such dependencies across hundreds of cancer cell lines (CCLs), forming the basis of the Cancer Dependency Map (DepMap)^20–22^. Integration of these datasets with molecular profiles of the screened models has enabled target and biomarker discovery^23^, and provides a foundation for developing predictive models of tumour vulnerabilities^24^.

However, identifying molecular features associated with dependency remains challenging because the number of candidate molecular features, across *omics*, exceeds the number of available CCLs by two or more orders of magnitude, creating a highly underdetermined, high-dimensional inference problem, prone to overfitting and spurious associations. In this context, machine-learning approaches can model complex relationships but sacrifice interpretability^25–27^, whereas simpler linear and logic models can identify more easily interpretable associations but require stringent prior feature reduction^28–30^. To tackle these challenges, we and others have previously proposed patient-guided feature selection, restricting dependency-association analyses to alterations observed in cancer patients^21,23,30^. However, because this strategy prioritises recurrent alterations, it biases discovery towards established oncogenic events and may overlook rare or weakly recurrent variants.

Here, we introduce CRISPR-VUS, a cancer-driver-agnostic framework that uses DepMap data to associate somatic variants with increased dependency on their host genes, without relying on patient recurrence or predicted functional impact during discovery. We call these variants Dependency-Associated Mutations (DAMs). Subsequently, we annotate DAMs with clinical recurrence, functional-impact predictions, druggability and drug-response associations to support interpretation and prioritisation.

We define DAMs that also associate with increased sensitivity to compounds targeting their host gene product as drug Sensitivity-Associated Mutations (SAMs). We then prospectively tested prioritised DAMs through genetic validation and exploratory pharmacological assays. Overall, CRISPR-VUS provides a systematic computational approach for identifying rare somatic variants linked to experimentally testable genetic dependencies and candidate therapeutic vulnerabilities.

The CRISPR-VUS Portal (https://vus-portal.fht.org/) enables interactive exploration of the complete DAM catalogue and its associated dependency, clinical and pharmacological annotations.

## Results

### Systematic identification of somatic mutations associated with genetic dependency

We analysed 977 CCLs spanning 36 cancer-types with somatic mutation and CRISPR dependency data^31–33^ (**Fig. 1a**, **Supplementary Table 1**, **Extended Data Fig. 1a** and **Methods**). We searched for variants associated with increased dependency on the gene harbouring the alteration, through cancer-type-specific analyses.

**Figure 1.**
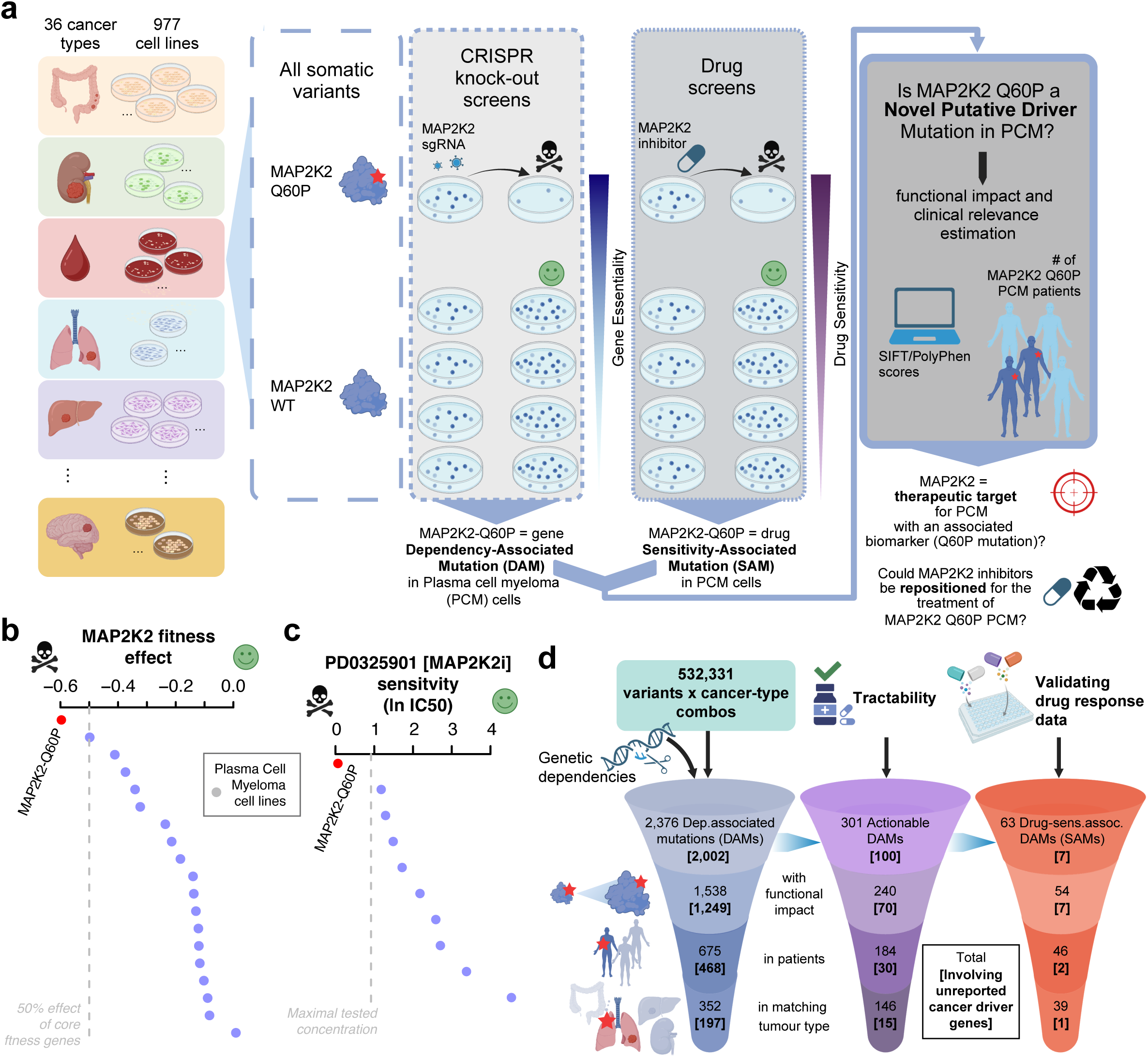
CRISPR-VUS pipeline and results overview. a. CRISPR-VUS integrates somatic-variant profiles with genome-scale CRISPR knockout data from 977 cancer cell lines (CCLs) across 36 cancer-types. Within each cancer-type, rankRatio tests whether CCLs carrying a variant rank among those most dependent on its host gene. Variants meeting predefined association, fitness-effect and empirical-significance criteria are classified as Dependency-Associated Mutations (DAMs). Integration with pharmacological screens identifies DAMs associated with increased sensitivity to compounds targeting the corresponding gene product, termed sensitivity-associated mutations (SAMs). Orthogonal annotations, including predicted functional impact, occurrence in patient tumours, histological concordance and target tractability, support interpretation and prioritisation. MAP2K2 p.Q60P in plasma cell myeloma illustrates the workflow. b. MAP2K2 CRISPR fitness effects across plasma cell myeloma CCLs. Dots denote CCLs; red denotes the MAP2K2 p.Q60P-bearing model. More negative values indicate stronger dependency. The dashed line indicates half the median effect of core-fitness genes. The variant-bearing CCL shows the strongest MAP2K2 dependency, thus MAP2K2 p.Q60P is classified as a DAM. c. Sensitivity of the same CCLs to the MEK1/2 inhibitor PD0325901, expressed as log of the half-maximal inhibitory concentration (IC50). The variant-bearing CCL (red) shows the lowest IC50. The dashed line indicates the maximal tested concentration. Its association with MAP2K2 dependency and PD0325901 sensitivity classifies MAP2K2 p.Q60P as a SAM. d. DAM possible prioritisation scheme using post-discovery annotations. Counts denote cancer-type-specific DAM instances; square brackets indicate subsets involving genes not currently classified as established cancer drivers.

We considered missense, in-frame insertion/deletion, and frameshift variants in genes not classified as common-essential or core-fitness genes^34^ (**Supplementary Dataset 1**). We selected these mutation classes because they can generate allele-specific functional effects^4,35^; we excluded other classes expected to cause generic loss of function, including nonsense, canonical splice-site and start-loss variants (**Methods**). We performed variant testing independently of cancer-driver annotation or predicted functional impact.

Most tested variants occurred in only one or a few CCLs (**Extended Data Fig. 1b**), making conventional association tests inapplicable or severely underpowered. To overcome this limitation, we developed the rankRatio score, which quantifies whether CCLs carrying a variant, or a set of variants within the same gene, rank among the most dependent on that gene within the cancer-type (**Fig. 1ab**; **Methods**). Importantly, the rankRatio evaluates singleton and recurrent variants within the same framework. Lower rankRatio values indicate stronger coupling between mutation status and dependency. For genes harbouring multiple variants across CCLs, the combination yielding the optimal rankRatio is retained (**Methods**).

DAMs were defined by a rankRatio < 1.71 (which is adaptively determined, **Methods**), median scaled CRISPR fitness effect < −0.5 (equivalent to half of the median fitness effect of core-fitness essential genes^34,36,37^) in mutant CCLs, and empirical permutation and hypergeometric probabilities below 20% (**Methods**).

We excluded genes carrying more than 10 distinct variants within each cancer-type-specific analysis to limit events compatible with loss of function or passenger accumulation in very long genes^13,17^.

We estimated a global false discovery rate (FDR) using variant instances occurring in genes not expressed in the mutant CCL as an empirical decoy population (**Methods**). At the rankRatio threshold of 1.71, the estimated FDR was 22%, remaining stable across the explored threshold range; the permutation and hypergeometric filters provided a modest improvement in specificity (**Extended Data Fig. 2a-c**, **Supplementary Dataset 2** and Supplementary Results).

We extended the rankRatio to pharmacological data, testing whether DAM-bearing CCLs were selectively sensitive to compounds targeting the gene product. DAMs satisfying this criterion were classified as drug-Sensitivity Associated Mutations (SAMs, **Fig. 1c**).

### An unbiased pan-cancer catalogue of Dependency-Associated Mutations

Across 36 cancer-types, we tested 532,331 variant/cancer-type pairs involving 403,269 unique variants in 16,842 genes. We identified 1,899 significant cancer-type-specific hits encompassing 2,376 DAMs, corresponding to 2,284 unique variants in 1,383 DAM-bearing genes (DAMbgs) (**Figs. 1d**, **2a**; **Supplementary Tables 2-3**). DAMs represented only 0.45% of all tested variant/cancer-type combinations.

**Figure 2.**
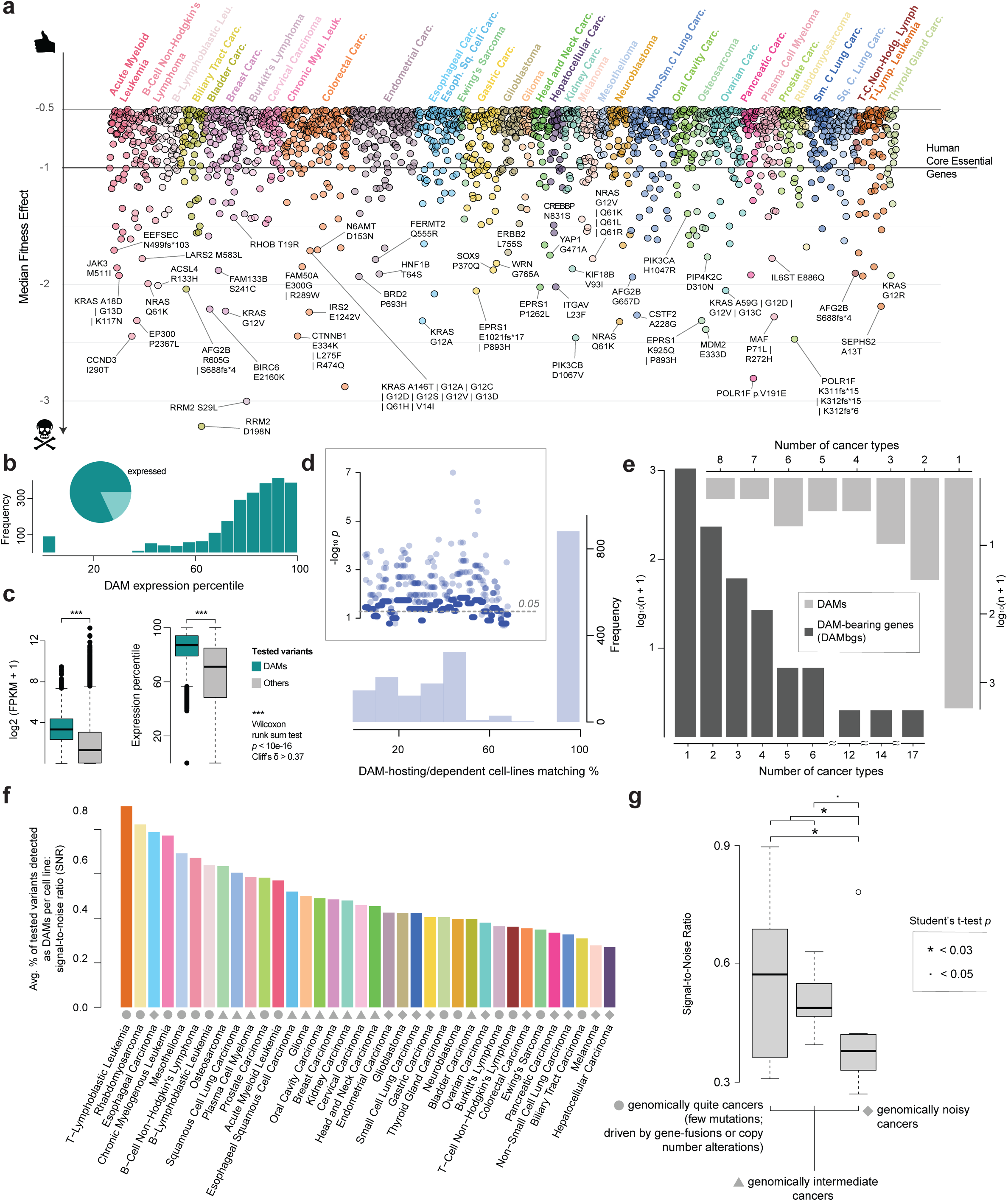
Landscape and biological properties of Dependency-Associated Mutations (DAMs) a. Cancer-type-specific hits, comprising individual DAMs or variant combinations meeting the rankRatio and fitness-effect criteria. Points denote hits and colours denote cancer-types. The y-axis shows the scaled fitness effect for individual DAMs or the median effect across mutant CCLs for combinations; all values are ≤ −0.5 with −1 marking the median effect of human core-essential genes. b. Proportion of DAMs in genes expressed in their host cancer cell line (CCL; FPKM > 1; pie) and distribution of within-CCL expression percentiles of DAM-bearing genes (histogram). c. Basal expression (log₂(FPKM + 1); left) and within-CCL expression percentile (right) of genes harbouring DAMs versus genes harbouring other tested variants in the variant-carrier CCLs. DAM-bearing genes show higher absolute and relative expression in the mutant CCL. Box plots show the median, interquartile range, whiskers extending to 1.5× the interquartile range and outliers. P values are from an unpaired Wilcoxon rank sum test with continuity correction (n = 585,344 variant instances). d. Distribution of concordance between CCLs hosting each hit and CCLs significantly dependent on the corresponding gene. Inset: Concordance significance as −log₁₀ P values from one-sided hypergeometric tests performed independently for each of the n = 1,899 cancer-type-specific hits; dashed line, P = 0.05. e. Numbers of DAMs (light grey) and DAM-bearing genes (dark grey) detected as hits in the indicated numbers of cancer-type-specific analyses. f. Signal-to-noise ratio (SNR), defined for each cancer-type as the average percentage of tested variants per CCL classified as DAMs. Colours denote cancer-types; symbols classify them as genomically quiet (circles), intermediate (triangles) or noisy (diamonds). g. SNR distributions across these genomic-complexity groups, defined from typical mutational burden and chromosomal instability. Individual cancer-types are observations. Box plots are defined as in c. Pairwise comparisons used paired Student’s t-tests (n = 36); * P < 0.03 and · P < 0.05.

Most DAMs were missense variants (90.9%), followed by frameshift (7.4%) and in-frame variants (1.7%)(**Extended Data Fig. 2d**). Frameshift DAMs displayed lower variant allele frequencies and less frequent loss of heterozygosity than background tested frameshifts (**Extended Data Fig. 2ef** and **Supplementary Results**). These observations are consistent with heterozygous contexts in which partial disruption of one allele may increase reliance on the remaining functional allele, analogous to previously described CYCLOPS dependencies^38^.

Eightytwo percent of DAM instances occurred in expressed genes, and they were generally enriched for occurrences in highly expressed genes (**Fig. 2bc**). Logistic regression further showed that the probability of DAM classification increased with the basal expression percentile of the host gene (**Extended Data Fig. 2g** and **Supplementary Results**). CCLs harbouring DAMs strongly overlapped with those showing dependency on the corresponding gene: 46% of hits showed perfect concordance and 78% significant overlap (**Fig. 2d**).

DAMs tended to be lineage-specific: 2,237/2,284 unique DAMs (98%) and 1,051/1,383 DAMbgs (76%) were identified in only one cancer-type-specific analysis (**Fig. 2e** and **Extended Data Fig. 1cd**). Among single-cancer-type DAMs, 73% (1,633/2,237) could not be tested elsewhere because the variant was absent from other cancer-types. Those testable in additional contexts showed little evidence of sub-threshold association (**Extended Data Figs. 3a**). At the gene level, 70.3% of single-cancer-type DAMbgs were represented by different variants in other cancer-types, whereas in 29.7% the same DAM-associated variant occurred elsewhere without generating dependency, supporting context-specific effects.

Although DAM numbers increased with the number of CCLs and tested variants, a CCL-wise signal-to-noise ratio normalised for mutation burden (**Methods**) showed the opposite trend: genomically quiet malignancies, including haematological cancers, sarcomas and prostate cancer, exhibited higher DAM-to-tested-variant ratios than highly mutated cancers such as colorectal cancer and melanoma (**Fig. 2fg**, **Extended Data Fig. 3b-d** and **Supplementary Results**). Thus, DAM discovery is not driven by mutation burden and enriches for highly expressed genes, missense variants and selective variant-dependency relationships.

### DAMs recover established cancer vulnerabilities and converge onto overlooked nodes in cancer-driver networks

DAMbgs significantly overlapped established cancer-driver catalogues (**Extended Data Fig. 4a**): 123 of 1,383 DAMbgs (8.9%) were classified as drivers in the IntOGen compendium^3^ (**Extended Data Fig. 4b**). This enrichment strengthened when considering only driver genes that were testable within our framework (**Fig. 3a**). For consistency, we used IntOGen membership as an operational benchmark throughout the study: we refer to these 123 genes as known DAMbgs and to the remaining 1,260 as unreported DAMbgs. The latter designation, however, does not imply an absence of previous experimental or literature evidence linking individual genes to cancer.

**Figure 3.**
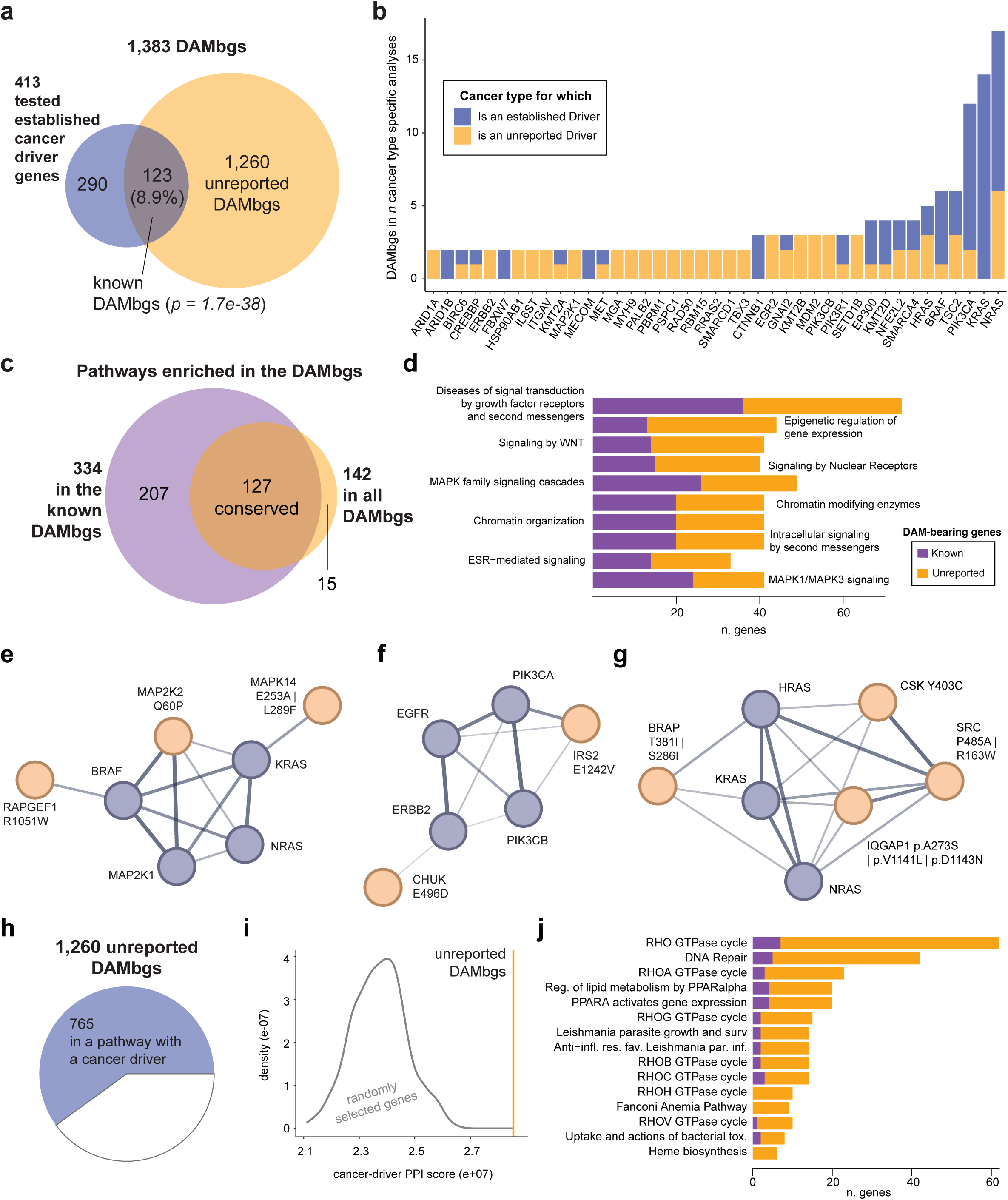
DAM-bearing genes characterisation and convergence on established cancer pathways. a. Overlap between 1,383 DAM-bearing genes (DAMbgs) and 413 tested established cancer-driver genes. Of the DAMbgs, 123 are established cancer drivers and 1,260 are unreported (one-sided hypergeometric test P = 1.7 × 10⁻³⁸). b. Known DAMbgs detected in at least two cancer-type-specific analyses. Each bar represents a gene; bar height indicates the number of analyses, and segments distinguish cancer-types for which the gene is already established (blue) or not (orange) as a cancer driver. c. Overlap between pathways enriched in known DAMbgs alone or in all the DAMbgs. Adding unreported DAMbgs preserved 127 enrichments (empirical P = 0.004) and yielded 15 additional enrichments (empirical P = 0.011). d. Ten most significant conserved pathway enrichments. Bar lengths indicate the number of DAMbgs; colours distinguish known (purple) and unreported (orange) DAMbgs. e-g. STRING interaction subnetworks illustrate DAMs in known and unreported DAMbgs within *Signalling to ERKs* (e), *PI3K/AKT Cascade* (f) and *Signaling by RAS mutants* (g). Purple nodes represent established cancer drivers and orange nodes represent unreported DAMbgs; edge widths are proportional to interaction confidence. h. Of 1,260 unreported DAMbgs, 765 (60.7%) share at least one biological pathway with at least one established cancer driver (empirical P = 2.38 × 10⁻⁷⁰). i. Cancer-driver protein-protein interaction (PPI) score for unreported DAMbgs (orange line) compared with the distribution obtained from randomly sampled, size-matched gene sets (grey curve). j. Pathways becoming significantly enriched only after adding unreported DAMbgs to the known DAMbgs. Bar lengths and colours are defined as in d.

Known DAMbgs were strongly enriched for genes harbouring activating mutations, consistent with the expectation that DAMs frequently capture oncogene addiction (**Extended Data Fig. 4ef**). Drivers including NRAS, KRAS and PIK3CA were recovered across cancer contexts (**Fig. 3b**). Conversely, analysed IntOGen drivers not recovered as DAMbgs were enriched for tumour suppressors or genes with ambiguous functional roles, supporting the specificity of the framework for dependency-generating alterations (**Extended Data Fig. 4cd** and **Supplementary Results**).

A total of 332 DAMbgs occurred in multiple cancer-types; 291 of them were unreported (**Fig. 3b**; **Extended Data Fig. 5a**). RIF1 exemplified recurrently nominated genes, with DAMs clustering in functionally relevant regions involved in replication timing and the 53BP1-RIF1 DNA-repair axis. Two RIF1 DAMs, p.E1620K and p.D495Y, are also observed in patient tumours matching the contexts in which they were identified in COSMIC^39^ (**Supplementary Table 5** and **Supplementary Results**).

We identified 187 cancer-type specific DAMs involving known DAMbgs but outside their recognised driver lineages (**Extended Data Fig. 5b**). Of these, 23 (12.3%) corresponded to canonical driver hotspots, including NRAS p.Q61K/L/R in B-cell non-Hodgkin lymphoma and bladder carcinoma, whereas 164 (87.7%) involved non-canonical variants (**Supplementary Table 3**). PIK3CA, for example, also emerged as DAMbg in chronic myelogenous and T-lymphoblastic leukaemia.

Multi-omic analysis provided little evidence that these observations resulted from CCL misclassification (**Extended Data Fig. 5c-h**), and DAMs were distributed across 596 CCLs rather than concentrated in a small number of models (**Extended Data Fig. 6a-c** and **Supplementary Results**).

At the pathway level, DAMbgs were enriched for established oncogenic signalling programmes, including MAPK, RAS, PI3K/AKT, VEGF, FGFR and PDGFR pathways (**Extended Data Fig. 7a** and **Supplementary Table 3**). Adding unreported DAMbgs to the known DAMbgs preserved 127 significant pathway enrichments, significantly more than expected by chance (**Fig. 3cd**, **Extended Data Fig. 7b** and **Extended Data Fig. 8a**, **Supplementary Table 3** and **Supplementary Results**).

Examples within ERK signalling include MAP2K2 p.Q60P, which we identified as a DAM in plasma cell myeloma (PCM), together with MAPK14 and RAPGEF1. Although activating MAP2K2 alterations have been reported in other tumour contexts^40,41^, MAP2K2 is not classified as a driver gene in the IntOGen benchmark, and p.Q60P has not been established as a driver alteration in PCM. Further examples include CHUK and IRS2 within PI3K/AKT signalling and BRAP, CSK, IQGAP1 and SRC within RAS-associated networks (**Fig. 3eg**).

A total of 765 unreported DAMbgs belong to at least one pathway alongside at least one established cancer driver gene significantly more frequently than expected by chance, and their protein-protein interaction connectivity to cancer drivers is significantly elevated (**Fig. 3hi** and **Extended Data Fig. 8de**).

15 pathways, including RHO GTPase signalling, DNA repair/Fanconi Anaemia and PPARα-associated metabolic programmes, emerged as significantly enriched only upon inclusion of the unreported DAMbgs (**Fig. 3j**, **Supplementary Table 3**). The likelihood of observing this, or a larger, number of new pathway enrichments by chance is 0.011 (**Extended Data Fig. 8b**, **Methods**), and the probability of identifying exactly these 15 pathways is less than 0.007 (**Extended Data Fig. 8c**, **Methods**). Established DNA-repair DAMbgs such as ATM, PALB2, POLQ, POT1 and RAD50 were already present; the addition of unreported genes including BRIP1, FANCD2, FANCI, FANCM, MRE11, MSH2, RIF1, WRN and XRCC4 expanded pathway coverage sufficiently for enrichment to reach statistical significance.

Together, these results show that unreported DAMbgs extended established oncogenic networks rather than creating unrelated pathway signals and that CRISPR-VUS identifies overlooked nodes within established cancer-driver networks.

### Unreported Dependency-Associated Mutations and Canonical-Oncogene-Addictions are largely mutually exclusive

We asked whether DAMs tend to emerge in CCLs that lack canonical oncogenic addictions. We classified a CCL as oncogene-addicted if it harboured a DAM in a gene annotated as a lineage-specific gain-of-function (GoF) driver. This definition is restrictive in requiring that the driver itself emerges as a DAM, but inclusive in that it does not distinguish between classical “addictive oncogenes” and other GoF drivers. By this criterion, 67% of CCLs with DAMs in unreported DAMbgs (n = 507) showed no co-occurring DAM in established drivers, with a median across cancer-types of 79% and complete absence of co-occurrence (100%) observed in 11 cancer-types (**Supplementary Table 4**, **Extended Data Fig. 6d**).

We also applied a more relaxed definition, considering a CCL oncogene-addicted if it harboured any lineage-specific GoF driver gene that was both mutated and essential, regardless of whether it was identified as a DAMbg. Even under this broader criterion, 21% of CCLs with DAMs in unreported DAMbgs lacked a mutated and essential oncogene, with a median of 18% across cancer-types and complete independence (100%) observed in three cancer-types (**Supplementary Table 4**, **Extended Data Fig. 6e**).

### Translational prioritisation and pharmacological vulnerabilities

We incorporated orthogonal annotations to prioritise DAMs. PolyPhen-2^42^ and SIFT^43^ predicted functional impact for 1,538/2,376 DAMs (65%) (**Extended Data Fig. 9a-d)**. DAMs were also enriched for clinically relevant CancerVar^14^ Tier I-II classifications (two-sided Fisher’s exact test, TSFET, P = 3.22×10⁻⁴⁸, n = 403,248), high REVEL^15^ scores (≥0.7; TSFET P = 0.005, n = 371,741), and elevated CADD^16^ scores (two-sided Mann-Whitney test P = 5.19×10⁻⁶, n = 403,220), providing independent support from existing variant-interpretation frameworks (**Supplementary Table 2**).

We assessed DAM recurrence in COSMIC^39^ and IntOGen^12^(**Extended Data Fig. 9ef)**. Overall, 1,032 DAMs were observed in patients, including 729 involving unreported DAMbgs (**Supplementary Table 5**). 675 DAMs were predicted to affect protein function and observed in patient cohorts. Of these, 352 occurred in patients with tumour types matching the CCL context in which the DAM was identified (**Fig. 1d**).

DAMs occurring in established cancer driver genes were the most frequently observed across patient cohorts (**Fig. 4a**, **Supplementary Tables 1** and **6**, **Extended Data Fig. 9h**). DAMs involving unreported DAMbgs were also recurrently detected in patients, often within the same cancer-type context in which they were identified (**Extended Data Fig. 9g** and **Supplementary Results**).

**Figure 4.**
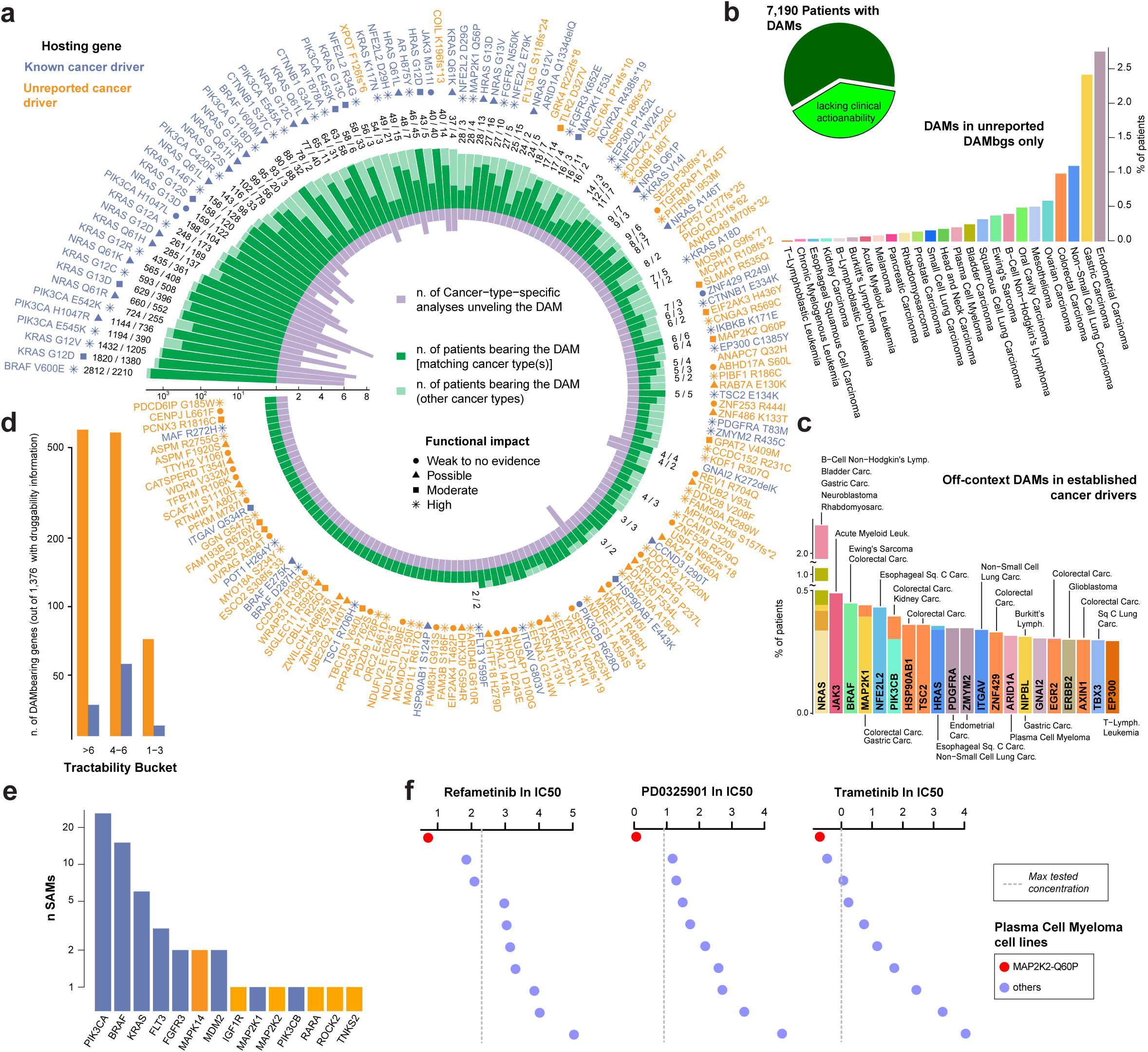
Patient prevalence, clinical actionability and pharmacological associations of Dependency-Associated Mutations (DAMs) a. Patient recurrence and cancer-type distribution of DAMs. Each sector represents a DAM observed in at least two patients whose tumour type matched at least one cancer-type in which the DAM was identified; variants are ordered by total patient prevalence. Outer labels indicate the affected gene and amino acid change and are coloured blue or orange according to whether the DAM-bearing gene is or is not an established cancer driver, respectively. On the logarithmic radial scale, light-green bars extend to the total number of patients carrying each DAM across all tumour types, whereas dark-green overlays indicate patients with tumour types matching those in which the DAM was identified. Adjacent numbers report total/lineage-matching patient counts. Inward purple bars show the number of cancer-type-specific analyses identifying each DAM. Symbols indicate functional-impact predictions derived from SIFT and PolyPhen: circle, weak or no evidence; triangle, possible; square, moderate; and asterisk, high. Absence of a symbol indicates unavailable predictions. b. Percentage of analysed COSMIC patients, by cancer-type, harbouring DAMs exclusively in unreported DAM-bearing genes and lacking actionable CIViC variants. The pie chart summarises this subgroup among 7,190 patients with DAMs. c. Percentage of patients harbouring off-context DAMs in established cancer-driver genes, identified in cancer-types not previously associated with the corresponding driver. Bars represent genes, and colours identify cancer-types. d. Distribution of 1,376 DAM-bearing genes with tractability information across tractability buckets. Colours distinguish established (blue) and unreported (orange) cancer-driver genes. e. Genes most frequently involved in sensitivity-associated mutations (SAMs). Bar height indicates the number of SAMs; colours are defined as in D. f. ln(IC50) values for refametinib, PD0325901 and trametinib across plasma cell myeloma CCLs. Red denotes the MAP2K2 p.Q60P-bearing CCL, blue denotes other CCLs with dashed lines indicating the maximal tested concentrations. MAP2K2 p.Q60P, a predicted functionally impactful DAM observed in multiple patients, consistently showed the lowest IC50 for all three MEK inhibitors, identifying it as a SAM and illustrating a potential drug-repurposing opportunity.

In COSMIC, 39% of patients harbouring a DAM lacked a co-occurring clinically actionable CIViC^44^ variant (**Fig. 4b, Extended Data Fig. 9ij**), suggesting that DAMs could contribute to stratification opportunities. Several DAMs in established cancer-driver genes were observed in patients with tumour types in which those genes are not currently recognised as drivers (**Fig. 4c**)

Among DAMbgs with druggability information^21,45^, 102 fell within tractability buckets 1-3, including targets of approved or preclinical agents (**Fig. 4d**, **Methods**). Integration of drug-response data from the GDSC^46^ identified 63 of them associated with sensitivity (SAMs) to 33 compounds (**Fig. 1d**, **Fig. 4e**, **Supplementary Table 2** and **Supplementary Results**). These included PIK3CA, KRAS, BRAF and FGFR3, as well as unreported DAMbgs.

MAP2K2-p.Q60P in plasma cell myeloma represented one case, associated with MAP2K2 dependency and sensitivity to multiple MEK1/2 inhibitors (**Fig. 1bc**, **Fig. 4f**).

### Independent computational convergence and prospective testing of prioritised DAMs

We compared our results with an independently conducted companion study by Sesia et al.^47^, developed in parallel and employing a distinct variant-processing pipeline. Sesia et al. restricted testing to recurrent variants while allowing associations beyond host genes.

Despite these and other analytical differences and a limited common background of only 6,768 tested variants (respectively 1.67% and 36% of all the tested variants in the respective studies), 65 DAMs overlapped significantly between the two analyses (21.8-fold enrichment; one-sided hypergeometric test P = 4.89 × 10⁻⁷⁸), including variants in five genes not classified as established cancer drivers (**Extended Data Fig. 10a-d** and **Supplementary Table 6**).

We next asked whether systematic CRISPR-VUS prioritisation could nominate rare variants with causal effects on cellular fitness.

We focused on non-small cell lung carcinoma (NSCLC) because an immortalised human alveolar type II epithelial model compatible with prime editing was available, enabling testing in a lineage-relevant background. Starting from 137 NSCLC-specific DAMs, we applied systematic annotation-based criteria requiring predicted functional impact, occurrence in at least two histology-matched patients and high-confidence target tractability. Only RTN4IP1 p.A80T and RHOT1 p.D243E satisfied these criteria (**Supplementary Table 2**, **Methods**). Here, tractability was used as a forward-looking target-prioritisation feature and did not require the availability of an existing selective inhibitor.

We tried to introduce these variants by prime editing into immortalised human alveolar type II epithelial cells with engineered *MLH1* loss to increase prime editing efficiency, and assessed cellular fitness through competition assays between variant-edited and wild-type cells (**Methods**). We successfully introduced RTN4IP1 p.A80T, whereas repeated attempts to generate RHOT1 p.D243E with three pegRNA designs did not yield detectable editing (**Extended Data Fig. 10e-f**).

Across three independent biological experiments under low-serum conditions, RTN4IP1 p.A80T conferred a significant competitive growth advantage relative to the non-targeting control, comparable to that produced by the oncogenic PIK3CA p.C420R DAM (**Supplementary Table 2**) included as positive control, with established activating allele^48^, available validated pegRNA construct and reproducible editing conditions (**Fig. 5a**, **Supplementary Table 7**). These results demonstrate that a rare DAM identified by CRISPR-VUS can prospectively generate a driver-like fitness phenotype in an independent epithelial model.

**Figure 5.**
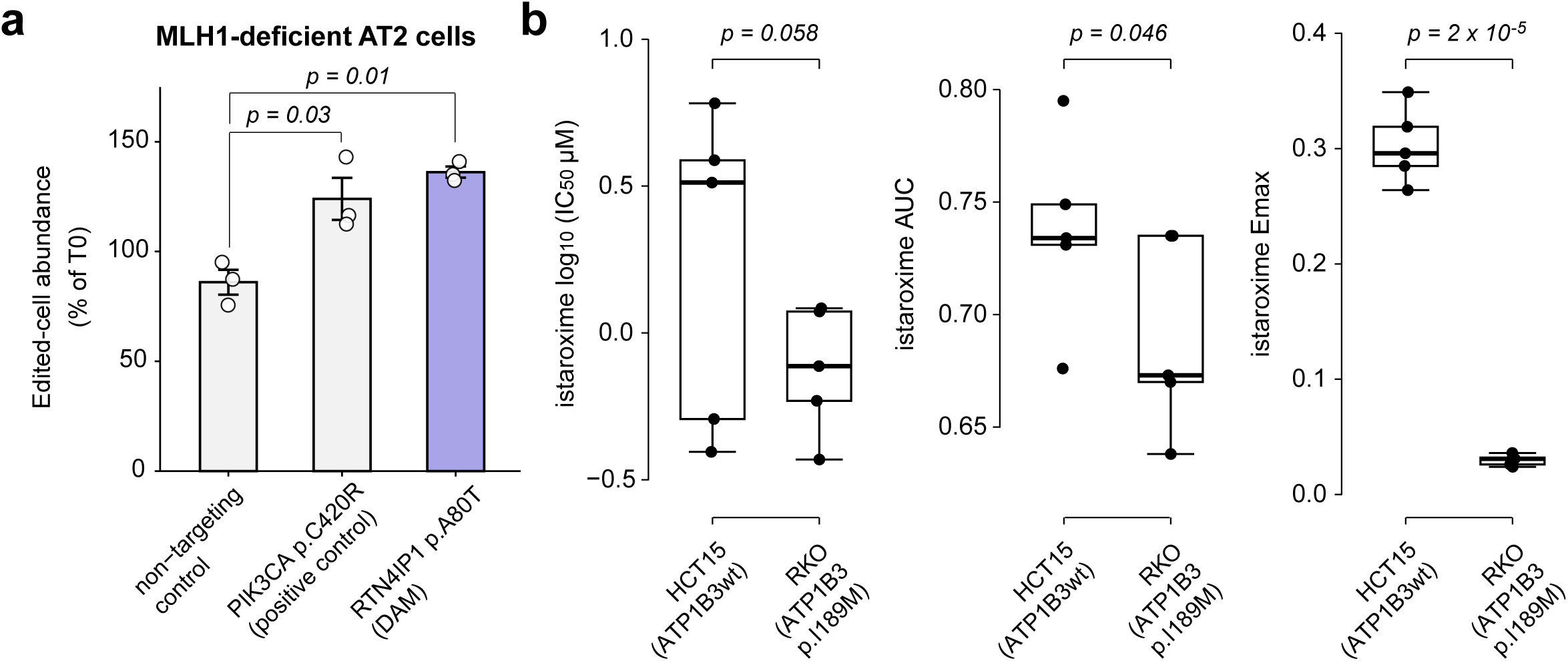
Prospective genetic and pharmacological validation of prioritised CRISPR-VUS predictions. a. Competitive-proliferation assays in MLH1-deficient, PE2-expressing human alveolar type II (AT2) cells carrying a non-targeting pegRNA, the PIK3CA p.C420R positive control or the RTN4IP1 p.A80T DAM. BFP-positive pegRNA-expressing cells were mixed 1:1 with GFP-positive parental cells and maintained under 0.5% serum; endpoint BFP:GFP ratios were normalised to time zero (T0). Points represent the mean of three technical wells from each of three independent experiments; bars show mean ± s.e.m. Differences were assessed by two-sided one-way repeated-measures ANOVA with experiment as the matching factor (F(2,4) = 14.39, P = 0.0149), followed by Dunnett’s comparisons with the non-targeting control (PIK3CA p.C420R, P = 0.03; RTN4IP1 p.A80T, P = 0.01). b. ATP1B3-wild-type HCT15 and ATP1B3 p.I189M-bearing RKO colorectal carcinoma cells were treated with istaroxime for 72 h under 0.5% serum. IC50, area under the dose–response curve (AUC) and fitted minimum relative viability (Emax) were estimated independently in five matched experiments, with four technical wells per concentration. Points represent independent experiments; boxes show the median and interquartile range, and whiskers extend to 1.5x the interquartile range. To test the predicted greater sensitivity in RKO cells, groups were compared using one-sided paired Student’s t-tests (log₁₀IC50, P = 0.058; AUC, P = 0.046; Emax, P = 2 × 10⁻⁵). Lower values indicate greater sensitivity. RKO and HCT15 are non-isogenic models; the analysis therefore assesses the reproducibility of the differential CCL response but does not isolate the causal contribution of ATP1B3 p.I189M.

Moreover, we considered the 15 DAMs at the final level of the actionable-DAM prioritisation funnel (**Fig. 1d**, **Supplementary Table 2**). These variants are predicted to affect protein function, observed in patients with matching tumour histology, hosted by genes not classified as established cancer drivers, and affect druggable proteins. Of these, we prioritised candidates that (i) could not previously be evaluated as SAMs because suitable drug-response data were unavailable in public screening repositories; (ii) had a predicted functional high-impact; (iii) were highly expressed and (iv) exhibited a scaled CRISPR fitness effect below −1 in the DAM-bearing cells, i.e. conferring a fitness loss that is at least as strong as the median effect observed for established essential genes, such as ribosomal protein genes. Because pharmacological perturbation generally produces less complete target inhibition than CRISPR knockout, we reasoned that such strong genetic dependencies would be more likely to produce a measurable differential drug response. Together, these criteria prioritised ATP1B3 p.I189M for experimental testing.

We performed an exploratory pharmacological test treating ATP1B3 p.I189M-bearing RKO colorectal carcinoma cells and ATP1B3-wild-type HCT15 cells with istaroxime, a drug under development for treatment of acute decompensated heart failure, targeting the Na⁺/K⁺-ATPase complex^49^, of which ATP1B3 encodes a β subunit^50^.

Across five independent paired experiments, RKO cells showed lower fitted residual viability at maximal response (Emax; one-sided paired Student’s t-tests, PST, P = 2 × 10⁻⁵, n = 10) and AUC (PST P = 0.046, n = 10), together with a non-significant trend towards lower log₁₀IC50 values (PST P = 0.058, n = 10; **Fig. 5b**, **Extended Data Fig. 10g** and **Supplementary Table 7**). Because this comparison involved two non-isogenic cell lines, variant status might be confounded with CCL identity. The experiment therefore demonstrates a reproducible difference in istaroxime response between RKO and HCT15 under the tested conditions but cannot establish that ATP1B3 p.I189M causes this difference or functions as a predictive biomarker.

Collectively, the prime-editing experiments establish a causal growth-promoting phenotype for RTN4IP1 p.A80T, whereas the istaroxime experiment provides preliminary CCL-level evidence consistent with the predicted ATP1B3-associated vulnerability. Isogenic models or broader panels of ATP1B3-mutant and wild-type cell lines will be required to test the variant-specific drug-response association.

## Discussion

CRISPR-VUS provides a framework for interpreting somatic variants by linking their occurrence to measured cancer-cell dependencies, i.e. Dependency-Associated Mutations (DAMs). By avoiding patient recurrence and prior driver annotation, the approach captures rare and singleton variants that are inaccessible to frequency-based cancer-driver analyses.

Several observations support the biological relevance of the catalogue. DAMs occur in expressed genes and recover oncogene addictions. A signal-to-noise analysis shows that genomically quieter cancer-types have a higher ratio of DAMs to tested variants than highly mutated cancers, consistent with dependency signals being less diluted by passenger-rich mutational backgrounds. DAMs converge on canonical cancer pathways, show connectivity to recognised cancer drivers and are enriched for variants predicted to be pathogenic or clinically relevant.

Their predominance in specific tumour contexts highlights the lineage dependence of genetic addiction.

Unreported DAMbgs extend rather than replace established cancer biology. Their convergence on MAPK, RAS, PI3K/AKT and DNA-repair programmes suggests that rare mutations can expose overlooked nodes within oncogenic networks. The absence of canonical oncogene addictions in many CCLs harbouring unreported DAMs indicates that some may define alternative survival mechanisms and vulnerabilities in poorly stratified tumours.

CRISPR-VUS provides a framework for translational prioritisation. Functional-impact predictions, patient recurrence, target tractability and drug-response associations are integrated after DAM discovery. This allows users to prioritise the catalogue according to biological or clinical objectives.

Introduction of the top-priority NSCLC-specific RTN4IP1 p.A80T DAM by prime editing conferred a reproducible competitive growth advantage in an independent, lineage-relevant epithelial model, demonstrating driver-like activity. Together with its association with selective RTN4IP1 dependency and evidence of future tractability, this finding nominates RTN4IP1 p.A80T as a previously unrecognised candidate oncogenic driver allele with the potential to define a therapeutically exploitable RTN4IP1 addiction in NSCLC. More broadly, this reinforces the potential of CRISPR-VUS to uncover rare oncogenic drivers and addictions, including dependencies involving druggable proteins, that may represent promising therapeutic targets and opportunities for drug repurposing.

Exploratory pharmacological testing showed a greater istaroxime response in RKO than in HCT15 cells, consistent with the predicted ATP1B3-associated vulnerability. However, because the two cell lines are non-isogenic and differ in their broader molecular backgrounds and growth properties, this experiment cannot attribute the response difference specifically to ATP1B3 p.I189M. Isogenic models or larger panels of mutant and wild-type cell lines will be required to determine whether the variant predicts istaroxime response.

Furthermore, CRISPR-VUS identifies associations rather than establishing causality, and individual predictions will require experimental validation in appropriate models.

Other limitations include incomplete representation of tumour subtypes in CCL collections and uneven availability of pharmacological data.

Future extensions could broaden CRISPR-VUS beyond coding sequence variants to regulatory alterations and copy-number changes, although these will require approaches to assign non-coding variants to context-specific target genes and to disentangle genuine copy-number-associated dependencies from regional and CRISPR associated biases^51^.

Coupling somatic variation with cancer dependency maps provides an orthogonal route to variant interpretation that complements recurrence- and annotation-based approaches. By identifying rare alterations associated with genetic addiction and pharmacological vulnerability, CRISPR-VUS expands the landscape of candidate cancer drivers, biomarkers and therapeutic opportunities beyond canonical oncogenic alterations.

The publicly available CRISPR-VUS Portal enables users to investigate whether variants detected in individual tumours have been associated with a dependency in lineage-matched cancer models and to prioritise these variants for further experimental assessment. If substantiated by experimental and clinical evidence, the catalogue could inform the design of focused sequencing panels, nominate candidates for prospective biomarker and patient-stratification studies, and prioritise druggable dependencies and drug-repurposing hypotheses for experimental assessment.

## Methods

### Input Data and the cancer cell lines (CCLs) analysis set

Full provenance information for all input datasets, including filenames, releases, URLs, download dates and content descriptions, is provided in **Supplementary Table 1**.

We obtained cancer cell-line (CCL) annotations, somatic variants, gene-expression profiles and gene-identifier mappings from the Cell Model Passports (CMP)^31^. CRISPR gene-effect scores were obtained from DepMap^20^ (24Q4), harmonised across the Broad and Sanger datasets, scaled as previously described^33,37^ and subjected to our previously established quality-control procedure^23^. Models absent from the CMP catalogue or assigned to non-specific or non-cancer categories were excluded. We analysed cancer-types represented by at least five eligible CCLs, resulting in 977 CCLs across 36 cancer-types (**Supplementary Table 1** and **Extended Data Fig. 1a**).

We obtained drug-response data from GDSC release 8.5^46^, cancer-driver annotations from IntOGen^52^ and additional benchmark catalogues^53^, target-tractability annotations from Open Targets^45^, patient somatic variants and sample annotations from COSMIC^39^, clinical-actionability annotations from CIViC^44^ and gene-level loss-of-heterozygosity calls from DepMap 24Q4. CMP model identifiers and approved HGNC symbols served as the primary model and gene identifiers, respectively. We harmonised cancer-type labels across resources using the manually curated mappings in **Supplementary Table 1**.

Additional dataset-processing and identifier-harmonisation details are provided in the **Supplementary Methods**.

### Selection of somatic variants for dependency testing

We performed the analysis separately within each cancer-type. Variants were grouped using the harmonised host-gene symbol and protein-level alteration. A variant observed in multiple cancer-types contributed one testable variant/cancer-type combination per cancer-type, whereas repeated observations within one cancer-type contributed multiple CCL instances to the same combination.

We excluded genes classified as core-fitness genes or common essentials^34^ because their knockout causes broadly deleterious effects independently of individual somatic alleles. We retained coding missense, in-frame insertion/deletion and frameshift variants, which can plausibly generate allele-specific functional consequences^4,35^. We excluded synonymous, nonsense, start-loss, stop-loss and essential splice-site variants. Within each cancer-type-specific analysis, we also excluded genes harbouring more than ten distinct qualifying variants to limit heterogeneous loss-of-function events and passenger accumulation in long genes.

Variant recurrence was not required: singleton variants were evaluated alongside recurrent variants. Cancer-driver status, predicted functional impact, patient recurrence, gene expression, tractability and pharmacological annotations were not used during DAM discovery and were incorporated only subsequently for interpretation and prioritisation.

After filtering, the discovery space comprised 532,331 variant/cancer-type combinations, corresponding to 403,269 distinct variants in 16,842 genes and 585,344 variant/CCL instances (**Fig. 1d**, **Supplementary Dataset 1**). Further details of variant-class selection are provided in the **Supplementary Methods**.

### Identification of Dependency-Associated Mutations (DAMs)

#### Cancer-type-specific dependency ranking and rankRatio calculation

We performed an analysis for each cancer-type individually, as follows. For each gene *g* with at least one observed variant across CCLs, we ranked all CCLs by decreasing dependency on *g* (i.e. increasing CRISPR-derived fitness effect). For a variant (or variant set) in *g* observed in *n* CCLs, we computed a rankRatio as the sum of the ranks of the *n* mutant CCLs divided by the Gauss’s sum at *n* (the sum of the first *n* natural numbers, or triangular number at *n*):

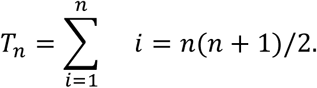

*T_n_* is the smallest possible rank sum. Consequently, the rankRatio is always greater than or equal to 1, with rankRatio = 1 when the *n* mutant CCLs occupy exactly the first *n* dependency ranks and, generally, with smaller rankRatio indicating stronger clustering of mutants among the most dependent CCLs.

For a singleton variant, *n* = *T_n_* = 1, thus the rankRatio is simply the dependency rank of its carrier. This allows singleton and recurrent variants to be evaluated within the same framework without requiring replicate mutant CCLs or estimating within-group variance.

#### Optimisation of variant combinations, rankRatio calibration and fitness-effect threshold

When multiple distinct variants of the same gene *g* were present across CCL from a given cancer-type, we applied an optimisation step: among all possible subsets of variants in *g*, we retained the one minimising rankRatio, ensuring that the dependency signal per gene/cancer-type pair reflected the most compelling association.

Because rankRatio has no parametric null distribution, we established a threshold empirically. The absence of a tractable null arises because rankRatio is a rank-based, ratio-normalised statistic that depends simultaneously on the number of mutant CCLs, their positions in the ranked dependency distribution, and the optimisation step when multiple variants per gene are present. As a result, the sampling distribution of rankRatio is non-standard: its variance and expected value change with *n*, the dependency score distribution within each cancer-type, and the number of possible variant subsets per gene. These dependencies prevent the derivation of a closed-form parametric null and render conventional significance models inappropriate, especially in the frequent case of singletons where no variance estimate is available.

To address this, we adopted an empirical benchmarking strategy using three archetypal oncogene addictions (BRAF in melanoma, KRAS in colorectal cancer, and PIK3CA in breast cancer). These dependencies were selected because they are well-established, lineage-specific genetic vulnerabilities with extensive experimental and clinical evidence supporting their functional relevance. We therefore reasoned that the least extreme rankRatio among these benchmark dependencies would provide a conservative empirical lower bound for defining biologically meaningful DAMs. This resulted in an operational rankRatio threshold of 1.71. To further enforce functional plausibility, we required that candidate variants display a median scaled fitness effect at least half the magnitude of that observed for known essential genes (e.g. ribosomal proteins).

#### Empirical permutation test and Hypergeometric rank-enrichment probability

For candidate hits passing the initial plausibility filters described above, we applied two orthogonal calibrations to test whether the observed clustering could plausibly arise by chance:

1. An empirical permutation P value (pₑₘₚ), obtained by shuffling dependency scores across CCLs within each cancer-type and recalculating rankRatio, mirroring the optimisation step.

Particularly, within each gene/cancer-type stratum, we permuted dependency scores across CCLs while preserving the marginal dependency distribution. For each permutation, we re-ranked the CCLs and recomputed rankRatio. Importantly, when multiple variants per gene were present, we re-applied the same optimisation step inside each permutation, selecting the variant subset with the lowest rankRatio. This symmetry ensures that the null distribution reflects the same search space as the observed data, thereby controlling for “winner’s curse”. This arises whenever one selects, among many candidate subsets, the one that optimises an objective function (here, the variant combination with minimal rankRatio). This selection process naturally biases the observed statistic downward relative to its expectation under the null. By mirroring the optimisation step within each permutation, we ensure that the null distribution is subject to the same bias, i.e. the null is also allowed to pick its best-performing subset. This symmetry prevents inflation of significance, making the resulting empirical P values conservative rather than anti-conservative.

The empirical one-sided P value was computed as:

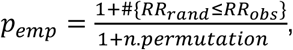

where *RR_rand_* is the rankRatio observed in a given permutation, *RR_obs_* is the observed one, # indicate the number of cases over all permutations and n. permutation = 1,000.

2. A one-sided hypergeometric tail probability quantifying the likelihood that n mutant CCLs would fall within the observed top-K ranks or higher under random placement.

To maximise robustness, we retained only candidate hits where both measures supported non-random clustering (P ≤ 0.2).

Each of these filters addresses a distinct dimension of plausibility: captures relative ranking, median fitness ensures consistency with dependency effect size, and permutation/hypergeometric P values provide statistical calibration against random expectations. These filters are thus layered rather than redundant, collectively enriching for variants most likely to reflect selective dependencies.

We deliberately adopted relatively lenient thresholds because the majority of DAMs are singleton or near-singleton events, where conventional multiple-testing frameworks lack power. Applying a universal false-discovery-rate (FDR) correction across hundreds of thousands of correlated tests would trivially discard nearly all signals. Furthermore, because our pipeline includes an optimisation step (evaluating multiple variant combinations per gene/cancer-type), the effective null space is neither uniform nor independent across tests, making global corrections conceptually inappropriate. Instead, we report empirical and hypergeometric P values transparently as complementary calibrations, while inference is strengthened by convergent orthogonal evidence including pathway and PPI enrichment, patient recurrence, functional impact predictions, signal-to-noise patterns, and overlap with independent studies.

#### Decoy-based estimation of global false discovery rate (FDR)

To complement the empirical permutation-based significance testing performed for each Dependency-Associated Mutation (DAM) or its optimal combinations with minimal rankRatio (therefore the HITS), we estimated the overall false discovery rate (FDR) of our discovery pipeline using an independent empirical decoy strategy.

As a conservative null set, we considered all variant occurrences in genes that were not expressed in the corresponding cancer cell-line (CCL). Variant occurrences in genes classified as non-expressed were assumed to be unlikely to generate genuine dependency-associated signals in that cellular context. Although we acknowledge that very lowly expressed genes may occasionally retain biological activity, we reasoned that this set provides a conservative approximation of the null distribution.

Across all analyses, we tested 532,331 variant/cancer-type combinations. Because the same variant can occur in multiple CCLs belonging to the same cancer-type, the total number of tested variant instances across all analyses was 585,344. Of these, 267,316 (46%) occurred in genes that were not expressed (FPKM < 1) in the corresponding CCL and therefore constituted the empirical decoy set (**Supplementary Dataset 2**).

We estimated the empirical global FDR across the full range of observed rankRatio thresholds, both before and after applying the HIT-local empirical and hypergeometric significance filters (P ≤ 0.2).

For a given rankRatio threshold value *τ*, we determined the corresponding HITS, defined as variants or combinations of variants with rankRatio ≤ *τ* and median scaled fitness effect of the mutated genes in the mutant CCL < −0.5 (corresponding to half the fitness effect reduction observed for core-fitness essential genes). The empirical permutation and hypergeometric significance filters were then optionally applied as described above.

We then grouped all individual variants across the HITS to obtain the set of unique Dependency-Associated Mutations (DAMs), while retaining each occurrence of a DAM in an individual cell line as a DAM-instance. The collection of DAM-instances therefore represents the complete set of positive predictions at a given rankRatio threshold *τ*.

At each threshold *τ*, the resulting DAM-instances were partitioned according to whether the mutated gene was expressed or not in the corresponding cell line. DAM instances occurring in non-expressed genes were considered putative false-positive predictions under our empirical null model.

Specifically, for each threshold *τ*, we computed:

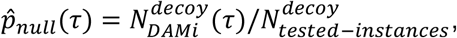

Where *N^decoy^*(*τ*) denotes the number of predicted DAM-instances (at the threshold *τ*) belonging to the empirical decoy set and *N^decoy^* is the total number of tested variant occurrences in non-expressed genes (267,316).

Under the assumption that the empirical null positive rate estimated from non-expressed genes is representative of the genome-wide background false-positive rate, we estimated the expected number of false-positive DAM-instances at threshold *τ* as:

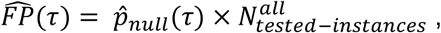

where *N^all^_tested-instances_* = 585,344.

Finally, we estimated the empirical global FDR as:

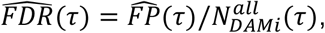

where *N^all^_DAMi_* (*τ*) is the total number of predicted DAM-instances at threshold *τ*.

This procedure was repeated for every *τ* between 1 and the operational threshold of 1.71, both before and after applying the empirical permutation and hypergeometric significance filters. The resulting FDR curves therefore quantify how the expected proportion of false-positive predictions varies with the rankRatio threshold and illustrate the contribution of the additional statistical filters to the overall specificity of the discovery pipeline. The estimated FDR remained stable across the range of rankRatio thresholds examined. At the operational threshold of 1.71, the estimated FDR was 22% after applying the empirical permutation and hypergeometric filters, compared with 23% without these filters. The filters reduced the total number of predicted DAM instances by 6% while reducing decoy DAM instances by 11%, indicating a modest but measurable improvement in specificity (**Extended Data Fig. 2a-c**, **Supplementary Dataset 2** and **Supplementary Results**).

#### Frameshift-DAM variant allele frequency and loss-of-heterozygosity analyses

Mutation consequence annotations and variant allele frequencies (VAFs) were obtained from the Cell Model Passports somatic-variant dataset described in the Input Data section.

Analyses were performed at the variant/CCL-instance level, because VAF and loss-of-heterozygosity (LOH) status are specific to the CCL carrying the variant. Accordingly, a variant observed in more than one CCL contributed one observation for each CCL. Instances lacking the information required for a given analysis were excluded without imputation.

For the VAF analysis, we compared frameshift DAM instances with missense and in-frame insertion/deletion DAM instances, considered jointly as the non-frameshift group. VAF distributions were compared using a two-sided Wilcoxon rank-sum test. We additionally calculated Cliff’s delta as a non-parametric measure of effect size, ordering the groups such that positive values indicated a tendency towards higher VAFs among missense and in-frame DAMs than among frameshift DAMs.

For the LOH analysis, we assigned each tested variant-CCL instance as LOH-positive or LOH-negative using binary data from the DepMap. We compared frameshift DAMs with the background of all other tested frameshift variants that passed the input-filtering criteria but were not classified as DAMs. We performed the corresponding comparison for missense and in-frame variants.

To determine whether the association between LOH and DAM classification differed according to mutation class, we fitted a binomial logistic-regression model with DAM status as the outcome and LOH status, mutation class and their interaction as explanatory variables. Mutation class was represented as frameshift versus missense/in-frame. The LOH-by-mutation-class interaction tested whether LOH distinguished DAMs from background variants differently for frameshift variants than for missense and in-frame variants. We also reported the proportion of LOH-positive instances within each group to facilitate interpretation.

We additionally fitted a binomial logistic-regression model with DAM status as the outcome and the within-CCL basal expression percentile of the host gene as the sole explanatory variable, at the variant/CCL-instance level, on n = 585,344 instances. Finally, we computed Odds ratios per 10-percentile increase.

All statistical tests reported in this section were two-sided and reported nominal P values. These analyses were performed after DAM identification and did not contribute to DAM calling or threshold selection.

#### Signal-to-noise ratio analysis

For each CCL, we calculated the percentage of tested variant instances that were identified as DAMs in the corresponding cancer-type-specific analysis. Variants belonging to an optimised multi-variant hit were counted individually as DAM instances. We then averaged these percentages across the CCLs belonging to each cancer-type to obtain the cancer-type-specific signal-to-noise ratio (SNR). Thus, higher SNR values indicate that a larger proportion of the variants tested in a cancer-type produced dependency-associated signals. We classified cancer-types as genomically quiet, intermediate or noisy based on their characteristic mutational burden and chromosomal instability. The classification assigned to each cancer-type is reported in **Supplementary Table 1**. We compared SNR values among these groups through unpaired Student’s t-tests.

#### Analysis of DAMs in established drivers outside their recognised cancer contexts, canonical-hotspot proportions

We classified DAM-bearing genes as established cancer drivers using the IntOGen^12^ compendium and defined their recognised cancer contexts from its cancer-type-specific annotations. Cancer-type labels were harmonised with the Cell Model Passports using the mappings in **Supplementary Table 1**. We classified a DAM/cancer-type association as off-context when IntOGen did not identify the corresponding gene as a driver in that cancer-type or an equivalent mapped category.

We quantified the prevalence of off-context DAMs in the corresponding COSMIC^39^ tumour cohorts by matching protein-level variant annotations. Patients carrying multiple qualifying DAMs in the same gene were counted once per gene/cancer-type combination.

To determine whether off-context DAMs corresponded to recognised driver alterations, we cross-referenced the 187 qualifying associations against the IntOGen driver-mutation catalogue. DAMs with an exact gene and amino-acid alteration match were classified as canonical driver mutations; all others were classified as non-canonical. Further details are provided in the **Supplementary Methods**.

#### Cell-line misclassification controls

We evaluated possible CCL misclassification using gene-expression, protein-abundance, CRISPR-dependency and drug-response profiles. For each CCL supporting an off-context DAM, we calculated Pearson correlations with all other eligible models and compared its similarity to its annotated cancer-type with its similarity to cancer-types in which the corresponding gene was an established driver. The target CCL was excluded from the reference set. Significance was assessed independently within each data modality by permuting cancer-type labels 1,000 times and calculating one-sided empirical P values. We considered evidence across all available modalities rather than relying on a single molecular profile. These post-discovery analyses did not alter CCL annotations or DAM classification. Further details are provided in the **Supplementary Methods**.

#### Identification of drug Sensitivity-Associated Mutations (SAMs)

We tested whether CCLs carrying a DAM also showed increased sensitivity to compounds targeting its host-gene product. DAMs satisfying this pharmacological association were classified as Sensitivity-Associated Mutations (SAMs).

DAM-bearing genes were linked to compounds using the curated drug–target associations from Gonçalves *et al.*^54^. Drug-response data were obtained from GDSC1 and GDSC2^46^ and represented as natural-log-transformed half-maximal inhibitory concentrations (ln(IC50)). A compound was considered eligible when the DAM-bearing gene was among its annotated targets, including for compounds with multiple targets.

Analyses were performed separately by cancer-type, starting from the 1,899 hits identified in the genetic-dependency analysis. For hits comprising an optimised combination of variants, the variant set selected during DAM discovery was kept fixed and was not re-optimised using drug-response data. Only CCLs with an available ln(IC50) for the compound under consideration were included.

For each compound and cancer-type, CCLs were ranked by ascending ln(IC50), with rank 1 assigned to the most sensitive model and tied values assigned their mean rank. We calculated a drug rankRatio (drug-RR) using the same rank-sum normalisation applied in the genetic analysis. A drug-RR of 1 indicates that the (n) DAM-bearing CCLs are the (n) most sensitive models. For singleton DAMs, drug-RR equals the sensitivity rank of the carrier.

We required drug-RR ≤ 1.5, a threshold calibrated using established pharmacogenomic associations in GDSC, including sensitivity of BRAF-mutant melanoma models to MAPK-pathway inhibition. We assessed non-random clustering using 1,000 one-sided permutations of ln(IC50) values among CCLs and the plus-one correction described above. Variant combinations remained fixed during permutation. We also calculated a hypergeometric tail probability, defining (K) as the rank of the least-sensitive DAM-bearing CCL and testing the probability that all (n) carriers occurred among the top (K) most sensitive models.

Hits with drug-RR <1.5 and empirical and hypergeometric probabilities ≤0.2 were classified as drug-sensitive. Their constituent variants were designated SAMs for the corresponding compound. We did not apply a global multiple-testing correction because this secondary analysis was restricted to previously identified DAMs and the available CCL sets differed among compounds and cancer-types. The resulting probabilities provide local calibration, and SAMs should be interpreted as candidate pharmacogenomic associations requiring independent validation.

#### Benchmarking against lists of established cancer drivers

We compared DAM-bearing genes (DAMbgs) with the IntOGen cancer-driver compendium and eight additional benchmark driver-gene lists compiled by Shi et al^53^ (**Supplementary Table 1**). Gene identifiers were harmonised to approved HGNC symbols, and each reference list was restricted to genes that were eligible for testing by CRISPR-VUS.

For each catalogue, we assessed the significance of the overlap with DAMbgs using a one-sided hypergeometric test, with all genes tested by CRISPR-VUS as the background.

IntOGen was used as the primary reference for classifying DAMbgs as established or unreported cancer drivers. Cancer-type-specific IntOGen annotations were used when evaluating whether the known DAMbgs occurred within or outside their previously recognised cancer contexts. These annotations were incorporated only after DAM discovery and did not influence DAM identification.

#### Functional Enrichment Analysis of the DAM-bearing genes

Reactome pathway-enrichment analysis was performed using the enrichPathway function from the ReactomePA R package^55^. Gene symbols were mapped to Entrez identifiers using biomaRt^56^. The background comprised all genes eligible for testing by CRISPR-VUS, excluding core-fitness genes and genes removed during variant filtering. Enrichment was evaluated separately for all DAM-bearing genes (DAMbgs), known DAMbgs listed in IntOGen and unreported DAMbgs. Pathways with a Benjamini-Hochberg-adjusted P value below 0.05 were considered significantly enriched.

To assess the contribution of unreported DAMbgs, we compared pathways enriched among known DAMbgs alone with those enriched after including all DAMbgs. Significance was evaluated using 1,000 Monte Carlo iterations in which the unreported DAMbgs were replaced by an equal number of randomly selected tested genes absent from IntOGen.

Sampling preserved the observed proportion of genes with and without Reactome annotations. At each iteration, enrichment analysis was repeated, and we recorded the numbers of conserved and newly significant pathways. Empirical P values were derived from the proportion of iterations producing at least the observed numbers of conserved or newly enriched pathways.

#### Protein-protein interaction network analysis of the DAM-bearing genes

Protein-protein interactions were obtained from STRING v12 using the STRINGdb R package^57^. We retained interactions with a STRING combined score of at least 200. The connectivity of unreported DAM-bearing genes (DAMbgs) to established IntOGen cancer drivers was quantified as the sum of the STRING edge weights linking genes across the two groups.

Statistical significance was assessed using 1,000 random gene sets, each matched in size to the unreported DAMbg set and sampled from genes represented in the STRING network after excluding established cancer drivers. For each random set, we recalculated its total connectivity to IntOGen drivers. A one-sided empirical P value was obtained from the proportion of random sets with connectivity equal to or greater than that observed for the unreported DAMbgs.

Illustrative pathway-specific subnetworks were extracted from the same STRING network, with edge widths proportional to the corresponding combined interaction scores.

#### Variant effect prediction analysis

We annotated variants using SIFT^43^ and PolyPhen-2^42^ predictions obtained through the Ensembl Variant Effect Predictor REST API using the GRCh38 assembly. When predictions were available for multiple transcripts, we retained the lowest SIFT score and highest PolyPhen-2 score.

Variants were assigned to five impact categories. Variants were classified as high impact when SIFT predicted a deleterious effect (score < 0.05) and PolyPhen-2 predicted a probably damaging effect (score ≥ 0.909). Moderate impact included variants predicted as deleterious by SIFT and possibly damaging by PolyPhen-2 (score 0.446-0.908), or as probably damaging by PolyPhen-2 alone. Variants supported as deleterious or possibly damaging by only one predictor were classified as possible impact. Variants with only tolerated or benign predictions were classified as low impact. Variants without applicable or available predictions, including non-missense variants, were classified as unknown impact.

These annotations were added after DAM discovery and did not contribute to DAM identification or threshold selection.

We additionally compared CRISPR-VUS classifications with CancerVar^14^, REVEL^15^ and CADD v1.7^16^. Analyses were performed on unique tested variants: variants identified as DAMs in at least one cancer-type were compared with variants never identified as DAMs. Missing scores were excluded separately for each tool without imputation.

For CancerVar, we compared the proportion of clinically relevant variants (Tiers I-II) with Tier III-IV variants using a two-sided Fisher’s exact test. REVEL score distributions were compared using a two-sided Mann–Whitney U test, and variants with scores ≥0.7 were additionally compared with lower-scoring variants using a two-sided Fisher’s exact test.

CADD PHRED-score distributions were analysed similarly; variants with scores ≥20, corresponding to the top 1% of predicted deleteriousness, were classified as high-CADD variants.

#### Tractability data and interpretation

We obtained gene-level tractability annotations from the Open Targets^45^ Platform dataset downloaded on 29 August 2025 and mapped targets to approved HGNC symbols. Where multiple modality-specific assessments were available, we retained the lowest-numbered, highest-confidence tractability bucket assigned to each gene.

As in our previous works^21,23^, we grouped genes into buckets 1-3, representing the strongest tractability evidence, buckets 4-6, representing intermediate experimental or structural evidence, and buckets above 6, representing more preliminary or indirect evidence. Lower-numbered buckets therefore indicate greater confidence that the encoded protein can be pharmacologically modulated. Higher-numbered buckets or missing annotations were not interpreted as evidence that a target is intrinsically undruggable.

Tractability annotations were incorporated only after DAM discovery and were used for translational interpretation and candidate prioritisation; they did not influence DAM identification.

#### Clinical relevance estimation of the DAMs

We assessed whether DAMs occurred in patient tumours using the COSMIC^39^ and IntOGen^12^ datasets described in the Input Data section. Variants were matched using harmonised gene symbols and protein-level alterations. Frameshift and deletion annotations were normalised to account for differences in notation between resources. When multiple nucleotide changes produced the same protein alteration, patients carrying any corresponding nucleotide-level variant were combined.

Cancer-type labels from COSMIC and IntOGen were mapped to the Cell Model Passports classifications using the manually curated mappings in **Supplementary Table 1**. A patient occurrence was considered tumour-type matched when the mapped tumour type corresponded to the cancer-type in which CRISPR-VUS identified the DAM. Cell Model Passports cancer-type definitions served as the reference classification.

Patient counts were de-duplicated within each resource. Multiple samples, sequencing records or matching nucleotide annotations from the same COSMIC patient were counted once per variant. IntOGen records were de-duplicated using cohort and patient identifiers. Because COSMIC and IntOGen do not provide a consistently shared patient identifier, counts were retained separately across the two resources rather than assumed to represent distinct patients. Patient prevalence was calculated relative to the number of unique analysed patients in the corresponding tumour cohort.

To identify patients lacking established actionable alterations, we used the CIViC Clinical Evidence Summaries^44^ described in the Input Data section. We retained sequence-level variants with clinical evidence and excluded molecular profiles representing fusions, expression changes, deletions, duplications, methylation, generic loss-of-function events or amplifications. CIViC disease labels were mapped to the Cell Model Passports cancer-types as specified in **Supplementary Table 1**. A patient was considered to harbour a clinically actionable alteration when at least one of their variants matched a retained CIViC entry in the corresponding tumour context. Patients carrying a tumour-type-matched DAM in an unreported DAMbg but no such CIViC alteration were classified as lacking a currently annotated actionable variant.

#### Co-occurrence analysis of DAMs and established driver mutations

We assessed co-occurrence at the CCL level, focusing on models harbouring at least one DAM in a gene absent from the IntOGen cancer-driver compendium. Each CCL was counted once, irrespective of the number of qualifying DAMs.

Under the primary definition, a CCL was classified as carrying a canonical oncogene addiction when it also harboured a DAM in a gene annotated by IntOGen as a gain-of-function driver for the corresponding cancer-type. As a sensitivity analysis, we applied a broader definition in which the CCL was required only to harbour a somatic mutation in a lineage-specific gain-of-function driver and to be dependent on that gene, defined by a scaled CRISPR fitness effect below −0.5, irrespective of whether the driver mutation was itself classified as a DAM.

For both definitions, we calculated the numbers and percentages of CCLs with unreported DAMs that did or did not harbour a co-occurring canonical dependency, both across the complete collection and separately by cancer-type. This analysis was descriptive and did not influence DAM identification.

#### Comparison of results with the independent study by Sesia et al

We compared our list of Dependency-Associated Mutations (DAMs) with the variants identified by Sesia et al.^47^ as positively associated with gene dependencies in CRISPR Cas9 screens. Because Sesia et al. performed pan cancer analyses that did not report tissue specific associations, we ignored cancer-type information in our comparison. We retrieved the set of variants shared between the two studies and used a one-sided hypergeometric test to assess the significance of their overlap, using as background all variants tested in both analyses.

#### Prioritisation of DAMs for prime-editing validation

We identified DAMs across four lung cancer categories: mesothelioma, non-small-cell lung carcinoma (NSCLC), small-cell lung carcinoma and squamous-cell lung carcinoma. We focused the experimental validation on NSCLC because an established immortalised human alveolar type II epithelial cell model suitable for prime editing and competitive-fitness assays was available. Alveolar type II cells provide a lineage-relevant cellular background for

NSCLC, particularly lung adenocarcinoma, whereas they are less representative of the mesothelial, neuroendocrine and squamous cellular origins of the other three cancer categories. This system therefore enabled selected NSCLC-specific DAMs to be introduced prospectively and their phenotypic effects evaluated in an independent, biologically relevant background.

Given the substantial effort required for prime editing and subsequent phenotypic characterisation, we applied a predefined, stringent data-driven strategy to prioritise candidates. Starting with 137 NSCLC-specific DAMs, we retained variants predicted to affect protein function, observed in at least two patients with matching NSCLC histology, and occurring in genes assigned to tractability Tier 1 (buckets 1-3) or the highest-confidence bucket of Tier 2 (bucket 4). Application of these criteria identified two candidates for experimental follow-up: RTN4IP1 p.A80T and RHOT1 p.D243E (**Supplementary Table 2**).

#### Prioritisation of DAMs for drug-screening validation

We considered the 15 DAMs at the final level of the actionable-DAM prioritisation funnel (**Fig. 1d, Supplementary Table 2**). These variants were predicted to affect protein function, occurred in patients with matching tumour histology, involved genes not classified as established cancer drivers and affected pharmacologically tractable proteins.

We prioritised candidates that had not been evaluated as SAMs because suitable cancer-type-matched drug-response data were unavailable in public screening repositories. We further required a scaled CRISPR fitness effect below −1 in the DAM-bearing CCLs, corresponding to a dependency at least as strong as the median effect observed for established essential genes. We selected this stringent threshold because pharmacological treatment generally produces less complete target inhibition than CRISPR knockout, making strong genetic dependencies more likely to generate measurable differential drug responses. Further imposing a “high impact” predicted functional impact and selecting the top expressed DAM-instance prioritised ATP1B3 p.I189M for experimental testing. Istaroxime was selected based on its pharmacological activity against the Na⁺/K⁺-ATPase complex^49^, of which ATP1B3 encodes a β subunit^50^. The availability of ATP1B3 p.I189M-bearing RKO cells and lineage-matched ATP1B3-wild-type HCT15 cells enabled direct comparative testing.

### Experimental validation

#### Cell culture

Immortalised (hTERT and SV40) human alveolar type II epithelial lung (AT2)^58^ cells were cultured in RPMI-1640 (Cat. No. 30-2001). The use of the AT2 cell line in this research was ethically reviewed by the Sanger REC and was granted a favourable opinion (S-REC 24_006). RKO and HCT15 colorectal carcinoma cells, harbouring the ATP1B3 p.I189M variant and wild-type ATP1B3, respectively, were maintained in ATCC-formulated EMEM and RPMI-1640, respectively. All media were supplemented with 10% FBS, 1% penicillin–streptomycin, and 1% L-glutamine, and cells were maintained at 37 °C in a humidified atmosphere containing 5% CO₂.

#### Generation of stable prime editing cell lines

To improve prime editing efficiency, AT2 cells were first made mismatch repair deficient through disruption of MLH1. MLH1 knockout (KO) was generated by transient transfection with a Cas9-T2A-EGFP plasmid (Addgene #48140) expressing an MLH1-targeting sgRNA (5′-GCACATCGAGAGCAAGCTCC-3′), as previously described^59^. Three days after transfection, GFP-positive single cells were isolated by fluorescence-activated cell sorting (FACS), expanded, and screened for MLH1 loss by western blotting.

The selected MLH1-deficient clone was subsequently engineered to express the prime editor through PiggyBac-mediated integration of a doxycycline-inducible PE2 expression cassette (PB-PE; Addgene #196971) together with the PiggyBac transposase plasmid, as previously described^60^. Cells were transfected with 1 μg of each plasmid using FuGENE HD (Promega) in Opti-MEM (Thermo Fisher Scientific). One week after transfection, GFP-positive single cells were isolated by FACS to establish PE2-expressing clones. GFP expression was routinely monitored as a surrogate marker of stable transgene integration, and prime editing activity was validated using the PE-FLARE reporter assay.

#### pegRNA design and cloning

pegRNAs targeting the RTN4IP1 A80T, RHOT1 p.D243E and PIK3CA C420R (used here as a positive control) variants were designed using PRIDICT 2.0^61^. The three highest-scoring pegRNA designs were experimentally evaluated, and the construct exhibiting the highest editing efficiency was selected for downstream validation experiments. Variant-targeting and non-targeting pegRNAs were cloned into a modified BbsI-digested pKLV2-hU6pegRNA(BbsI-ccdB)-PGK-Puro-T2A-BFP vector (Addgene #67974) by Gibson Assembly (New England Biolabs) to generate enhanced pegRNAs with a *tevopreQ1* hairpin. All pegRNA sequences are provided in **Supplementary Table 7**.

#### Prime editing validation and competitive proliferation assays

Stable AT2-PE2 MLH1-KO cells were transduced with pegRNA constructs targeting RTN4IP1 A80T, PIK3CA C420R, or a non-targeting control. 48 hours post-transduction cells were selected in growth medium (RPMI supplemented with 10% FBS, 1X penicillin-streptomycin) containing 2 µg/ml puromycin (Invivogen) for 4 days to generate stable lines. Following seven days of prime editing induced with doxycycline treatment (1 µg/ml), successful installation of endogenous edits was confirmed by Sanger sequencing (Eurofins). Primer sequences are listed in **Supplementary Table 7.**

To determine the effect of each variant on cellular fitness, competitive proliferation assays were performed by mixing BFP-positive, pegRNA-expressing/non-targeting cells with GFP-positive parental PE2-expressing cells at a 1:1 ratio immediately before plating. Cells were seeded at 500,000 cells per well in a 6-well plate and maintained under reduced-serum conditions (RPMI supplemented with 0.5% FBS) to enhance the detection of fitness phenotypes. At the indicated time points, cells were harvested by trypsinisation, washed in FACS buffer (PBS containing 0.5% FBS and 2 mM EDTA), stained with Zombie Red viability dye (BioLegend), and filtered through a 40-μm cell strainer before analysis.

Changes in the relative abundance of BFP-positive and GFP-positive populations were quantified by flow cytometry using LSRFortessa (BD Biosciences). BFP:GFP ratios were normalised to the initial ratio measured immediately after plating (day 0). An increase in the proportion of BFP-positive cells over time was interpreted as a proliferative advantage conferred by the edited variant. Each experiment was performed in technical triplicate and on three separate days. Technical replicates were averaged within each experiment. Endpoint-normalised BFP:GFP ratios were analysed by one-way repeated-measures ANOVA, with experiment as the matching factor, followed by Dunnett’s multiple-comparisons test comparing each edited condition with the non-targeting control. All tests were two-sided.

#### Drug dose-response assay

RKO and HCT15 cells were seeded in white, optical-bottom 96-well tissue-culture plates (Cat. No. 165306; Thermo Fisher Scientific) at densities of 2.1 × 10³ and 3.2 × 10³ cells per well, respectively, in 100 µL of medium. After 24 h, the medium was replaced with 100 µL of medium containing 0.5% FBS and istaroxime hydrochloride (Cat. No. HY-15718A; MedChemExpress). Cells were treated with eight concentrations of istaroxime, ranging from 0.00001 to 100 µM in tenfold increments, with four technical replicate wells per concentration. Vehicle-control wells received 0.1% DMSO.

After 72 h of treatment, cell viability was measured using the CellTiter-Glo 2.0 Luminescent Cell Viability Assay (Cat. No. G9243; Promega). Luminescence was measured using a Spark multimode microplate reader (Tecan). Signals from cell-free wells were subtracted as background, and viability was expressed relative to the corresponding vehicle-treated controls.

Five independent experiments were performed, with RKO and HCT15 cells tested in parallel within each experiment. Dose–response analyses were performed in RStudio using the GRmetrics package^62^. Curves were fitted separately for each cell line and experiment, and log₁₀-transformed half-maximal inhibitory concentration (log₁₀IC50), area under the dose– response curve (AUC) and fitted minimum relative viability (Emax) were calculated.

Technical replicates contributed to curve fitting but were not treated as independent observations.

The resulting metrics were compared between cell lines using one-sided paired Student’s *t*-tests, with independent experiment as the pairing factor. Pairing accounted for the fact that the two cell lines were tested concurrently under the same experimental conditions. We used directional one-sided paired tests because CRISPR-VUS predicted greater sensitivity in ATP1B3 p.I189M-bearing RKO cells, corresponding to lower log₁₀IC50, AUC and Emax values. An effect in the opposite direction would therefore not have supported the predicted pharmacological vulnerability. Statistical tests were based on five independent paired experiments.

#### The CRISPR-VUS Web Portal

The CRISPR-VUS Web Portal (https://vus-portal.fht.org) enables interactive exploration of the results reported in this study. Users can browse or search for genes and variants and examine their DAM status across cancer-types. Gene-specific heatmaps display patient prevalence, tumour context and predicted functional impact from SIFT and PolyPhen-2.

Selecting a DAM provides access to the underlying CCL-level dependency data and, where available, drug-sensitivity profiles, including IC50 values and tested concentration ranges.

The portal was implemented as a Vue.js single-page application with D3.js visualisations, a FastAPI backend and a relational MySQL database.

#### Plots and Figures

Plots were generated in R 4.4.2, illustrations in **Fig. 1** were created using BioRender.com.

#### Use of language-assistance tools

We used ChatGPT (OpenAI, GPT-5) as an advanced grammar and style assistant to improve clarity and readability of the manuscript. The tool was not used to generate, modify, or interpret scientific content, analyses, or results.

## Supplementary Material

**Supplementary Table 1** - Full provenance information for all input datasets; Fully annotated List of 977 cancer cell lines included in the analysis; Cross-resource cancer-type annotation harmonised mapping; list of established cancer driver genes from IntOGen and other state-of-the-art repositories; cancer-type genomic-noise groups.

**Supplementary Table 2** - Full lists of Dependency-Associated Mutations (DAMs) or their optimal combinations (HITS), individual cancer-type-specific DAMs, DAMs instances, DAM instances decoupled, and drug Sensitivity-Associated Mutations with scores and prioritisation criteria; DAM tissue-specificity summary; DAM functional impact predictions.

**Supplementary Table 3** - Functional characterisation of the DAM-bearing genes (DAMbgs); pathway enrichment analysis and convergence of the unreported DAMbgs onto established cancer driver networks; off-context DAM and hotspots analysis.

**Supplementary Table 4** - Co-occurrence of unreported DAMs with DAMs in known DAMbgs or established cancer drivers with activating mutations that are also essential.

**Supplementary Table 5** - Number of patients across cohorts and resources, harbouring each DAM; DAM co-occurrence in patients with clinically actionable mutations (according to CIViC); data underlying the sunburst plot in Fig. 4a.

**Supplementary Table 6** - Results from benchmarking Dependency-Associated Mutations with variant hits reported in Sesia et al.

**Supplementary Table 7** - Results from the experimental validation of prioritised DAMs by prime-editing CRISPR screen and pharmacological inhibition, underlying Fig. 5.

**Supplementary Dataset 1** - Full list of tested variants. Available on FigShare at: https://doi.org/10.6084/m9.figshare.33435853

**Supplementary Dataset 2** - Full list of tested variant instances with decoy set specification. Available on FigShare at: https://doi.org/10.6084/m9.figshare.33435853

## Supplementary Reusults

### Supplementary Methods

#### Data availability

All datasets and results are available within the Supplementary Tables, on FigShare (at https://doi.org/10.6084/m9.figshare.33435853) or through the external resources and portals specified in **Supplementary Table 1** and in the **Methods**. Processed data, including the full list of Dependency-Associated Mutations (DAMs), Sensitivity-Associated Mutations (SAMs), their annotations, and occurrences in patient cohorts can be accessed via the CRISPR-VUS Portal (https://vus-portal.fht.org).

#### Code availability

The complete computational workflow is available at https://github.com/francescojm/CRISPR-VUS. The repository contains a Jupyter notebook, accompanying R scripts, configuration files, software-environment information and instructions for reproducing the analyses and figures. The notebook can also be accessed through Google Colab (at https://colab.research.google.com/drive/1PkduxjbHQH4zZxk3g69LWQUGL_jsL-aq?usp=sharing).

Required input datasets are available on Figshare at https://doi.org/10.6084/m9.figshare.33435853 (CRISPR-VUS-DataPackage.zip) with the directory structure described in the code repository README file.

## Supporting information

Supplementary Results

## Acknowledgements

We thank Livio Trusolino, Andrea Bertotti, Emanuel Gonçalves and Mathew Garnett for critically reading the manuscript and providing valuable feedback.

We acknowledge the services provided by the National Facility for Data Handling and Analysis, IU3 – Technology Development - DevOps and Web Development Unit, Fondazione Human Technopole, Milan, Italy, and particularly thank Francesca Bonato for her contributions to the development, optimisation and maintenance of the CRISPR-VUS Portal.

## Extended Data Figure Legends

**Extended Data Fig. 1.**
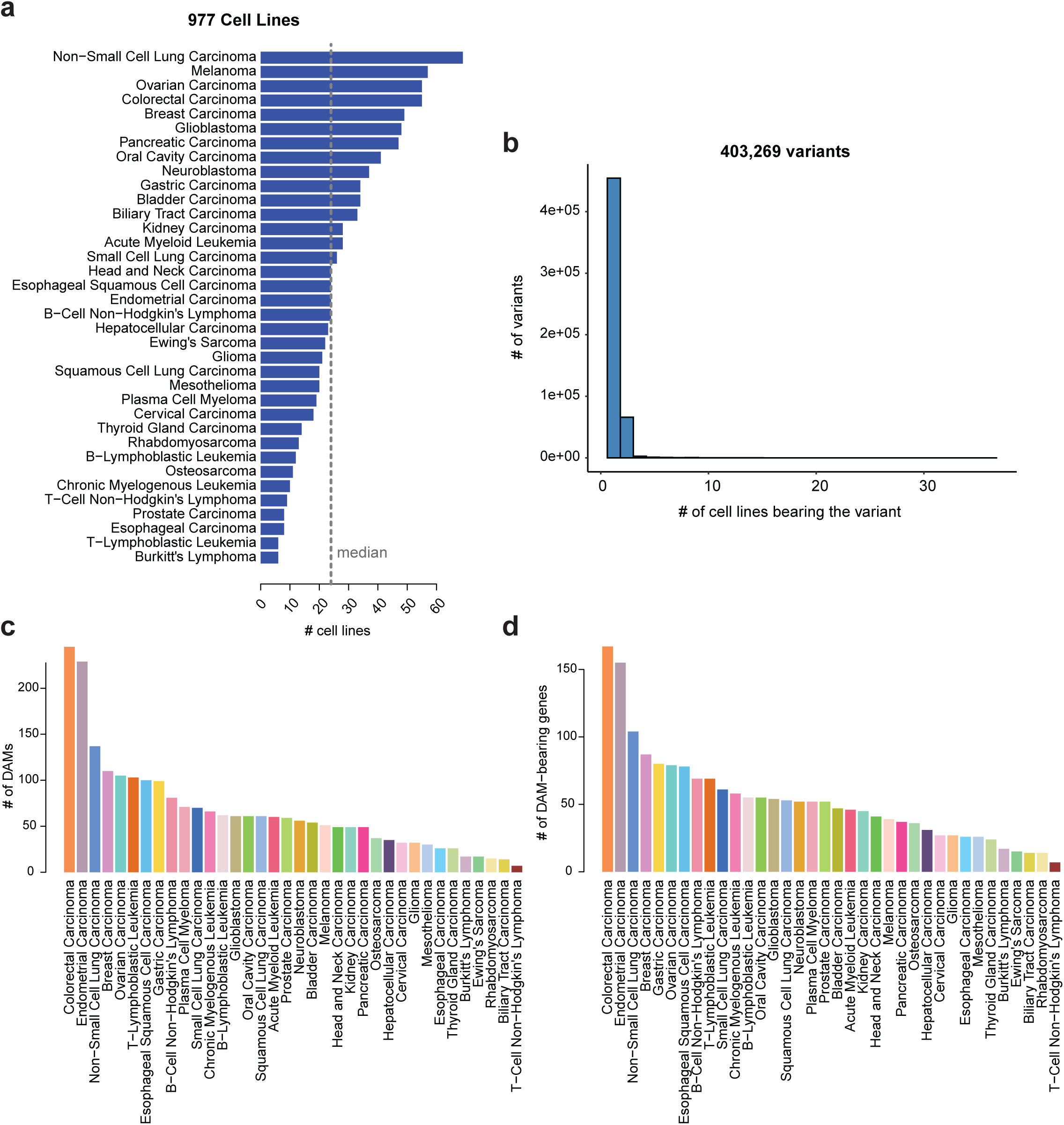
a. Cancer cell lines included in the analysis across cancer types. b. Number of tested variants across the number of carrier cell lines included in the analysis. c. Number of Dependency-Associated Mutations (DAMs) across cancer-type-specific analyses. d. Number of DAM-bearing genes across cancer-type-specific analyses.

**Extended Data Fig. 2.**
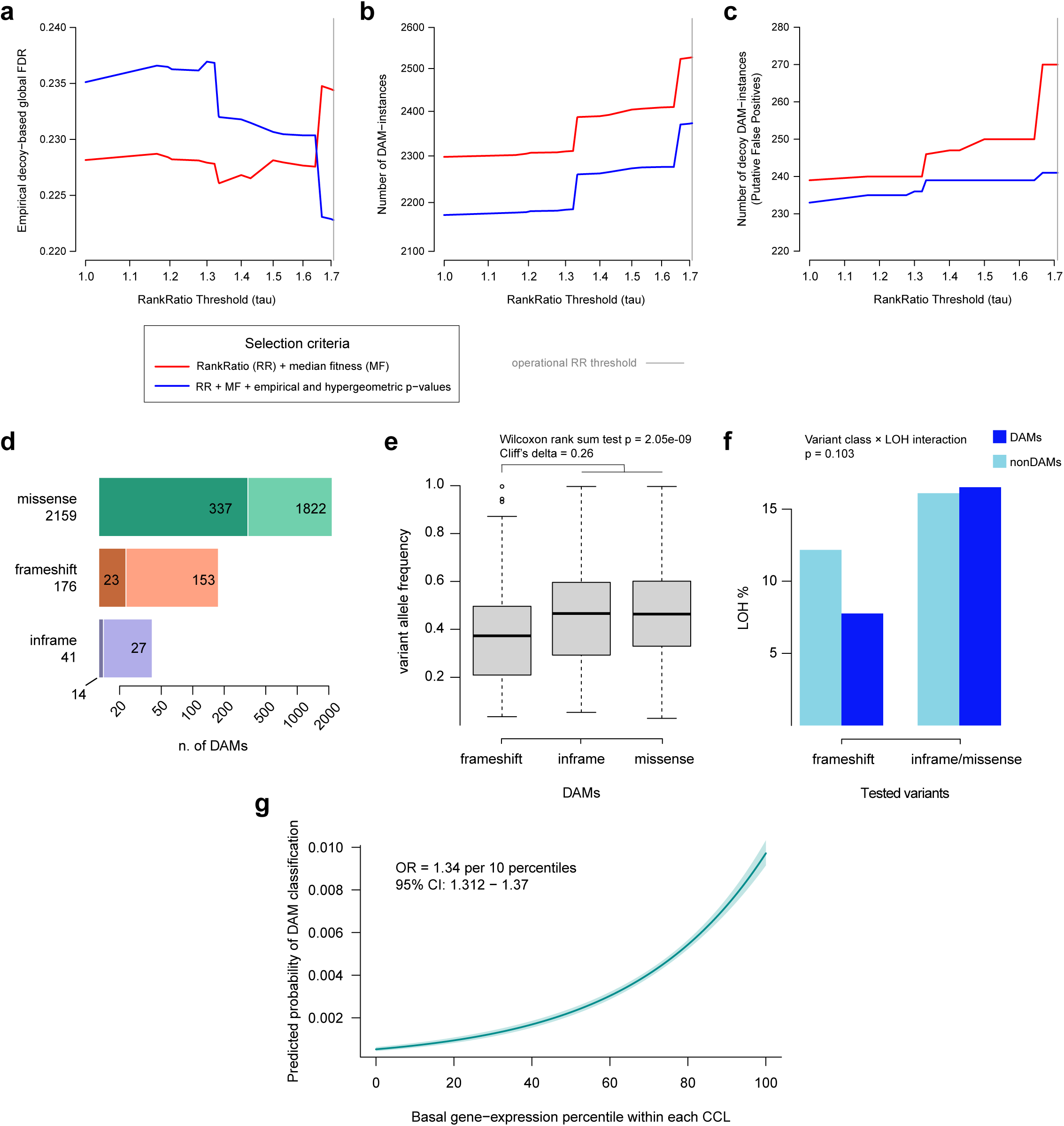
a-c. Empirical decoy-based estimation of the global false discovery rate of the CRISPR-VUS framework. a. Estimated empirical global false discovery rate (FDR) as a function of the rankRatio threshold (tau). b. Total number of predicted Dependency-Associated Mutation instances (DAM instances) across rankRatio threshold values. c. Number of predicted DAM instances belonging to the empirical decoy set, defined as variant instances occurring in genes that are not expressed (FPKM < 1) in the corresponding carrier cell line and used as putative false-positives under the empirical null model. Red curves correspond to DAM selection based on rankRatio (RR) and median fitness (MF) criteria alone, whereas blue curves additionally require empirical permutation and hypergeometric significance (P < 0.2). The vertical grey line indicates the operational rankRatio threshold (tau = 1.71) adopted throughout the study. A global FDR was estimated using the empirical decoy strategy described in the Supplementary Methods. d-f. Functional classes and variant allele frequencies of DAMs. d. Distribution of the 2,376 identified DAMs according to mutation class. Missense variants constitute the vast majority of the catalogue (90.9%), followed by frameshift (7.4%) and in-frame insertion/deletion variants (1.7%). Within each mutation class, darker and lighter shades indicate DAMs occurring in established cancer driver genes and in genes not previously classified as cancer drivers, respectively. Numbers within each segment indicate the corresponding number of DAMs. e. Distribution of variant allele frequencies (VAFs) across the three mutation classes. Frameshift DAMs exhibit significantly lower VAFs than the combined set of missense and in-frame DAMs, consistent with an enrichment of heterozygous events among frameshift DAMs. Boxes represent the interquartile range (IQR), centre lines denote the median, whiskers extend to 1.5 × IQR, and points indicate outliers. f. Percentage of variants occurring in genomic regions exhibiting loss of heterozygosity (LOH) among tested variants (light blue) and Dependency-Associated Mutations (DAMs, dark blue), shown separately for frameshift variants and for the combined set of missense and in-frame variants. Frameshift DAMs exhibit a lower proportion of LOH than the background distribution of tested frameshift variants, whereas the LOH frequency remains largely unchanged for missense and in-frame variants. g. Basal gene expression predicts the probability of identifying a DAM. Logistic regression relating the probability that a tested variant instance was classified as a DAM to the basal expression percentile of its host gene within the corresponding cancer cell line (CCL). The solid line represents the fitted logistic model, and the shaded region indicates the 95% confidence interval. Every 10-percentile-point increase in gene expression was associated with a 34% increase in the odds of DAM classification (odds ratio = 1.34, 95% CI 1.312-1.37, P < 9.95 × 10⁻¹⁵²).

**Extended Data Fig. 3.**
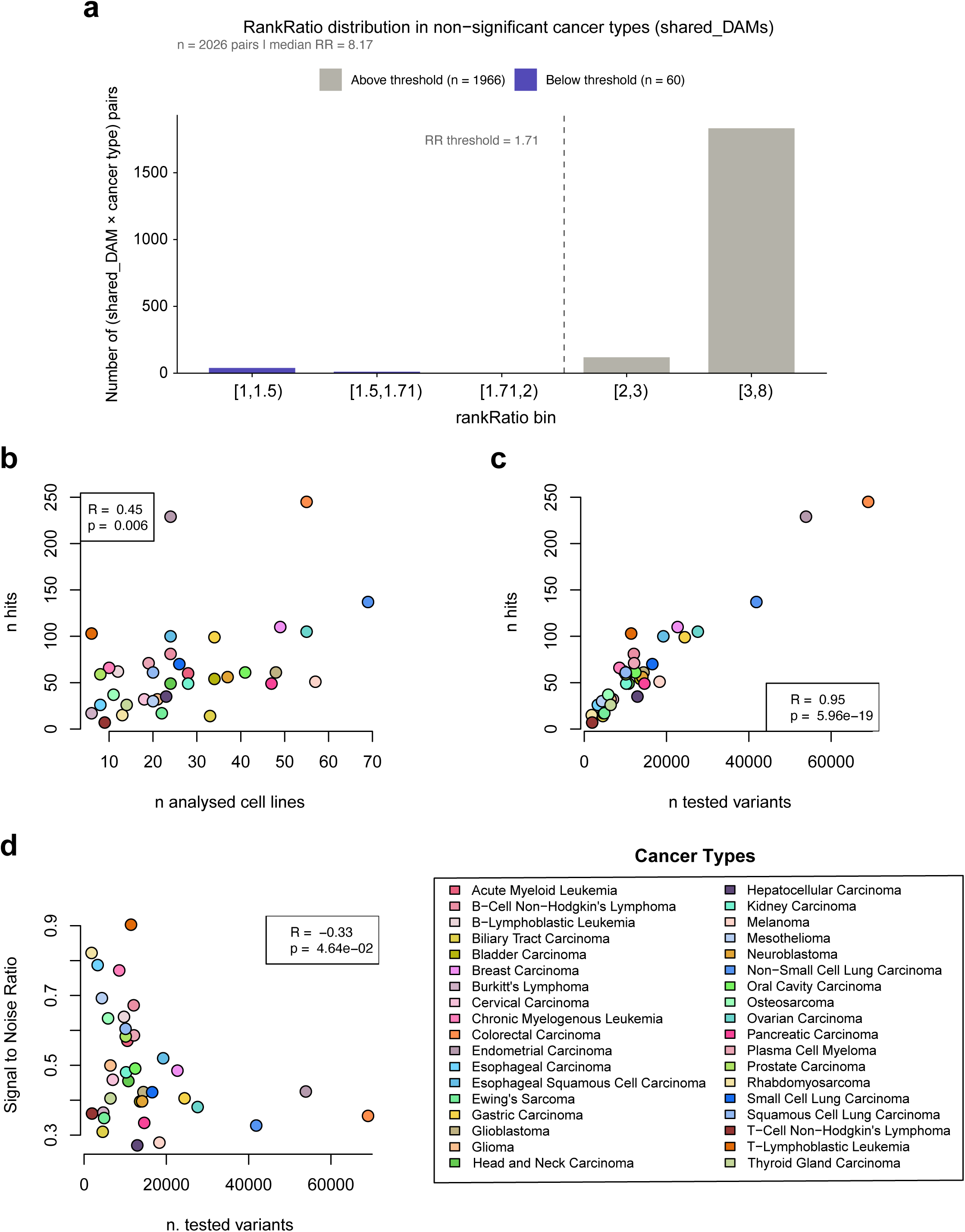
a. rankRatio distribution for shared-DAMs in non-significant cancer types. For each shared-DAM (a variant testable in ≥2 cancer types but significant in exactly one), the rankRatio (RR) is extracted from all cancer types where it was tested but did not reach significance. The figure shows the distribution of 2,026 (shared-DAM × cancer type) pairs, binned by RR value. Bars coloured in purple represent pairs falling below the pipeline RR threshold of 1.71 (the dashed line); grey bars indicate pairs above the threshold. b. Number of cell lines included in the analysis (x-axis) and number of hits (y-axis), i.e DAMs or combinations of them with optimal rankRatio and median fitness effect < -0.5, across cancer-types (the individual points) as indicated by the different colours. The values in the insets indicate Pearson’s correlation between the variables on the two axes and Pearson’s correlation test p-value, respectively. c. As in b., but with the number of tested variants on the x-axis. d. As in b and c, but with the number of tested variants on the x-axis and Signal-to-Noise Ratio (i.e. average number of DAMs per cell line divided by the average number of tested variants per cell line) on the y-axis.

**Extended Data Fig. 4.**
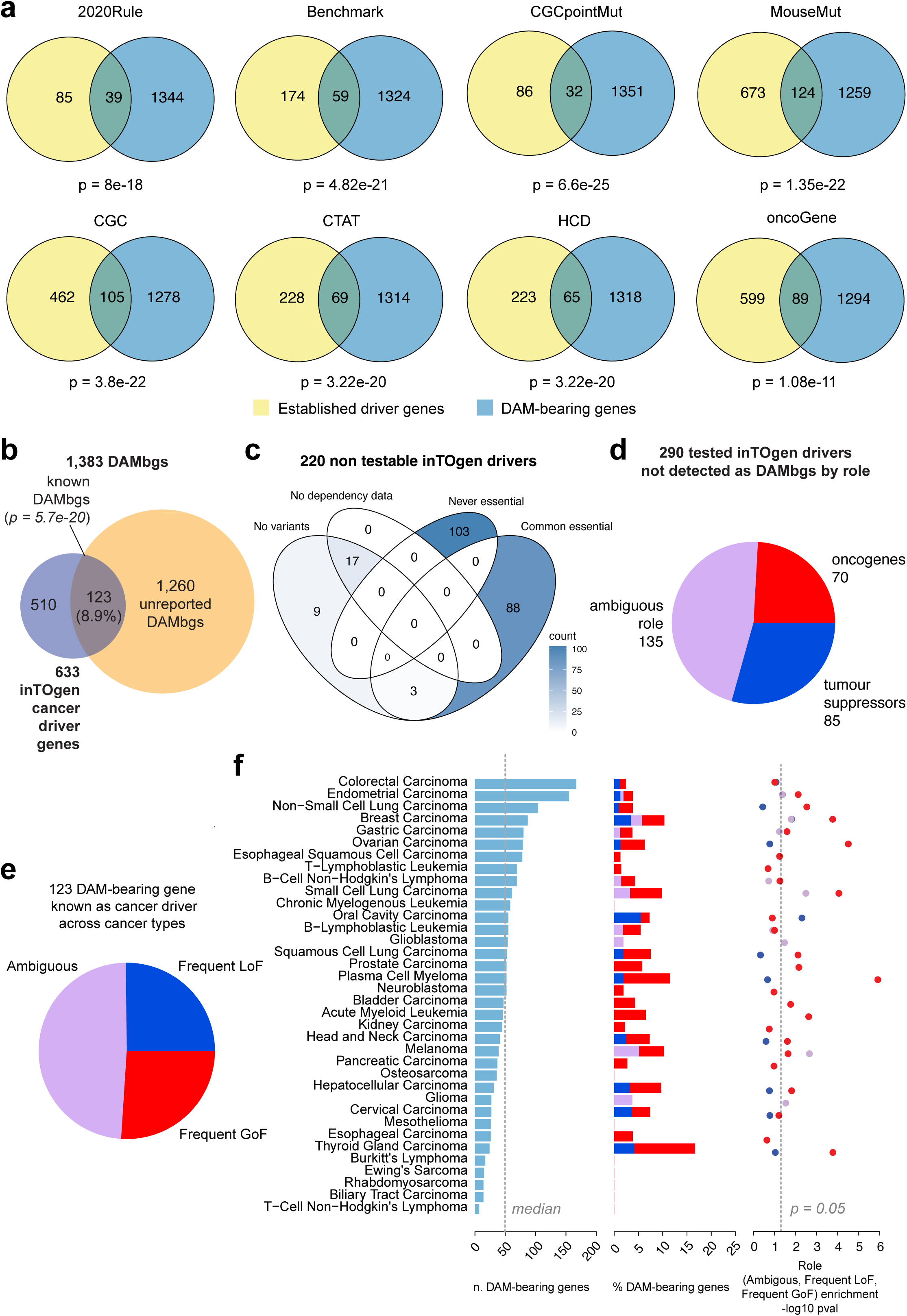
a. Overlaps between the DAM-bearing genes (DAMbgs) and lists of established cancer driver genes from multiple sources (Methods). The overlap is statistically significant (Fisher’s exact test P-values are reported) for all the considered sources. b. Enrichment of cancer driver genes annotated in the IntOGen census in the DAMbgs. This analysis uses the complete set of known IntOGen cancer driver genes as the reference background and therefore reflects the overall performance of the CRISPR-VUS framework, including both the analytical method and the limitations imposed by the underlying mutation and dependency datasets. Of the 633 IntOGen cancer driver genes, 123 (19.4%) are identified as DAMbgs (hypergeometric test, P = 5.7 × 10⁻²⁰). c. This panel refers, instead, to the enrichment analysis restricted to the subset of IntOGen cancer driver genes that were actually amenable to analysis (Fig. 3a), illustrating the reasons for the non-inclusion of 220 IntOGen drivers. Among the 510 driver genes not identified as DAM-bearing genes in panel b., 220 could not be evaluated at all because they lacked suitable somatic variants in the analysed cell lines (“No variants”), lacked CRISPR dependency data (“No dependency data”), were never sufficiently essential to satisfy the framework inclusion criteria (“Never essential”), or corresponded to common/core-essential genes excluded a priori from the analysis (“Common essential”). After excluding these non-testable genes, the effective background consists of 413 analysable cancer driver genes, of which 123 (29.8%) were recovered as DAM-bearing genes, resulting in an even stronger enrichment (hypergeometric test, P = 1.7 × 10⁻³⁸). d. Functional classification of the 290 tested IntOGen cancer driver genes that were not identified as DAMbgs. Only 70 (24%) are annotated as unambiguous oncogenes, whereas the remainder correspond to tumour suppressors (n = 85) or genes with context-dependent or ambiguous functional roles (n = 135). e. DAMbgs, annotated as cancer driver (according to IntOGen, known DAM-bearing genes) split based on their annotated cumulative role across cancer-types: frequently harbouring loss-of-function mutations (LoF), frequently harbouring gain-of-function mutations (GoF) or both (Ambiguous). f. All DAMbgs, across cancer-types (left plot), are split based on their annotated cancer-type-specific role (central plot). On the rightmost plot: Hypergeometric test enrichment p-values of each annotated role in the known DAM-bearing genes across cancer-types. Driver genes frequently harbouring GoF mutations are more prominently enriched than LoF and Ambiguous in the DAM-bearing genes across cancer-types.

**Extended Data Fig. 5.**
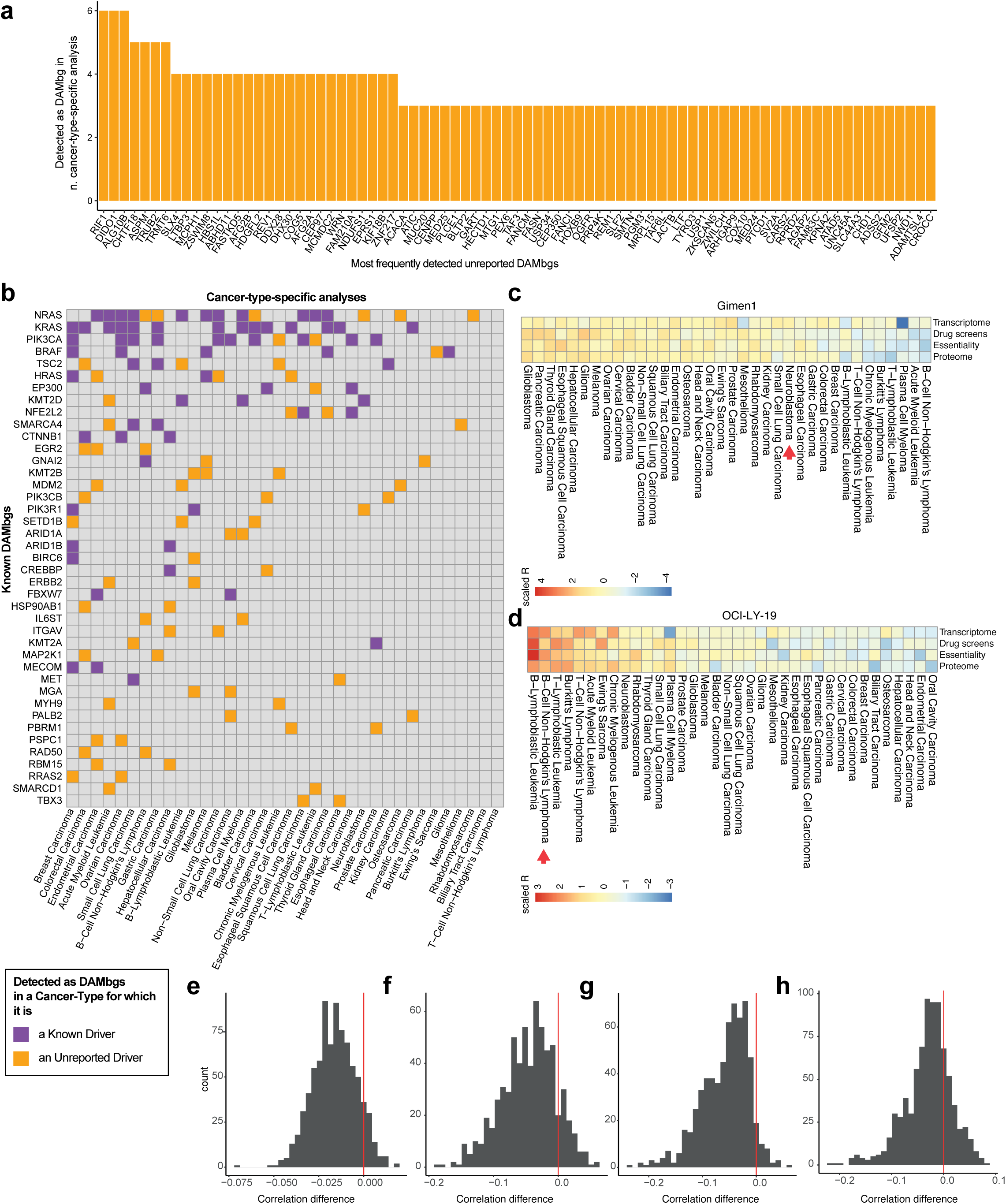
a. Unreported DAM-bearing genes (DAMbgs) most frequently detected across cancer-type-specific analyses. b. Known DAMbgs (on the rows) detected in at least two cancer-type-specific analyses (on the columns). Colours indicate whether the gene in the row was detected as DAMbg in a cancer-type for which it is known to be a cancer-type-specific driver. c-d. Lack of evidence of widespread cell line misannotations. c. Gimen1 (example of a cell line carrying an off-context DAM) correlation with cell lines belonging to each analysed cancer-type, based on transcriptomic, drug screens’ IC50s, essentiality and proteomic profiles. The red arrow indicates the original label. d. OCI-LY-19 (another example) correlation with cell lines belonging to each analysed cancer-type, based on transcriptomic, drug screens’ IC50s, essentiality and proteomic profiles. The red arrow indicates the original label. e-h. For each CCL harbouring an off-context DAM, we computed its mean Pearson correlation with the models of its own annotated cancer type (excluding itself) and with those of the alternative cancer type, i.e. the one in which the DAM-bearing gene is an established IntOGen driver. Panels show the distribution of the difference (alternative minus own) across all CCL–gene–alternative-cancer-type combinations tested, using CRISPR essentiality (e), drug IC50 (f), proteomic (g) and transcriptomic (h) profiles. Negative values indicate that the CCL resembles its annotated lineage more closely; positive values, expected under misclassification, indicate the opposite. The red line marking zero on the x-axis is included for reference.

**Extended Data Fig. 6.**
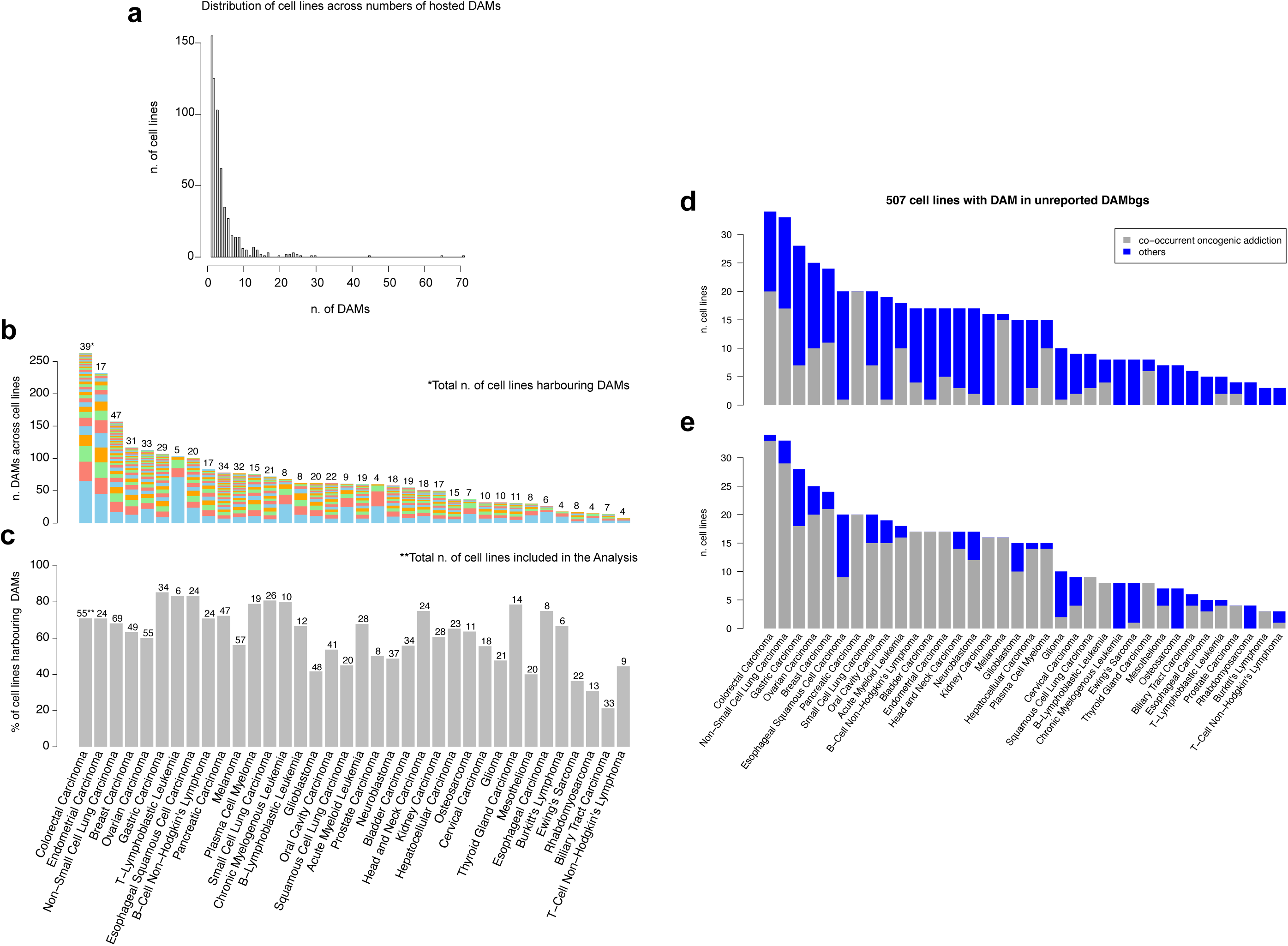
a. The distribution of the number of cancer cell lines (CCLs, y-axis) harbouring a fixed number of Dependency-Associated Mutations (DAMs, x-axis) shows most CCLs harbour one or a few DAMs. b. Number of observed DAMs across CCLs across cancer-types. Colours indicate different CCLs within cancer types, and the number of CCLs with at least one DAM for each cancer-type is indicated on top of each bar. c. Percentages of analysed CCLs harbouring at least one DAM, across cancer-types, with numbers indicating the total number of CCLs included in the analysis. d. Number of CCLs across analysed cancer-types that harbour DAMs in unreported DAM-bearing genes (DAMbgs) and are (grey bars) or are not (blue bars) also addicted to an established oncogene. Two criteria are used to define oncogenic addicted CCLs, respectively for the two plots. e. A CCL is oncogene-addicted if it harbours a DAM in a cancer-type-specific cancer driver frequently harbouring activating mutations. b. A CCL is oncogene-addicted if it harbours a mutation in a cancer-type-specific cancer driver gene that is also essential in that CCL.

**Extended Data Fig. 7.**
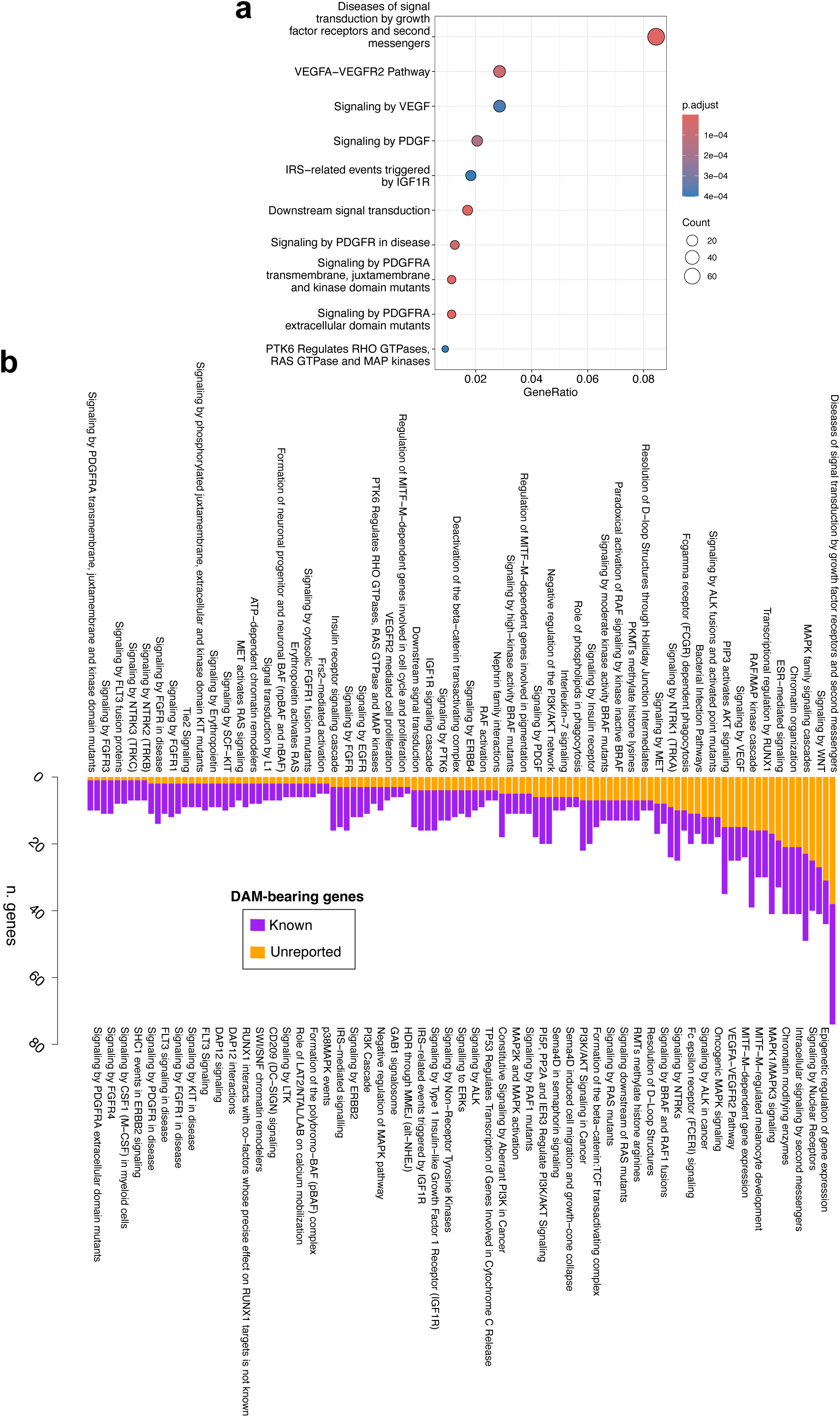
a. Top ten enriched Reactome pathways in the DAM-bearing genes. b. Reactome pathway enrichments in the known DAMbgs that are conserved when adding the unreported DAMbgs to the enrichment analysis.

**Extended Data Fig. 8.**
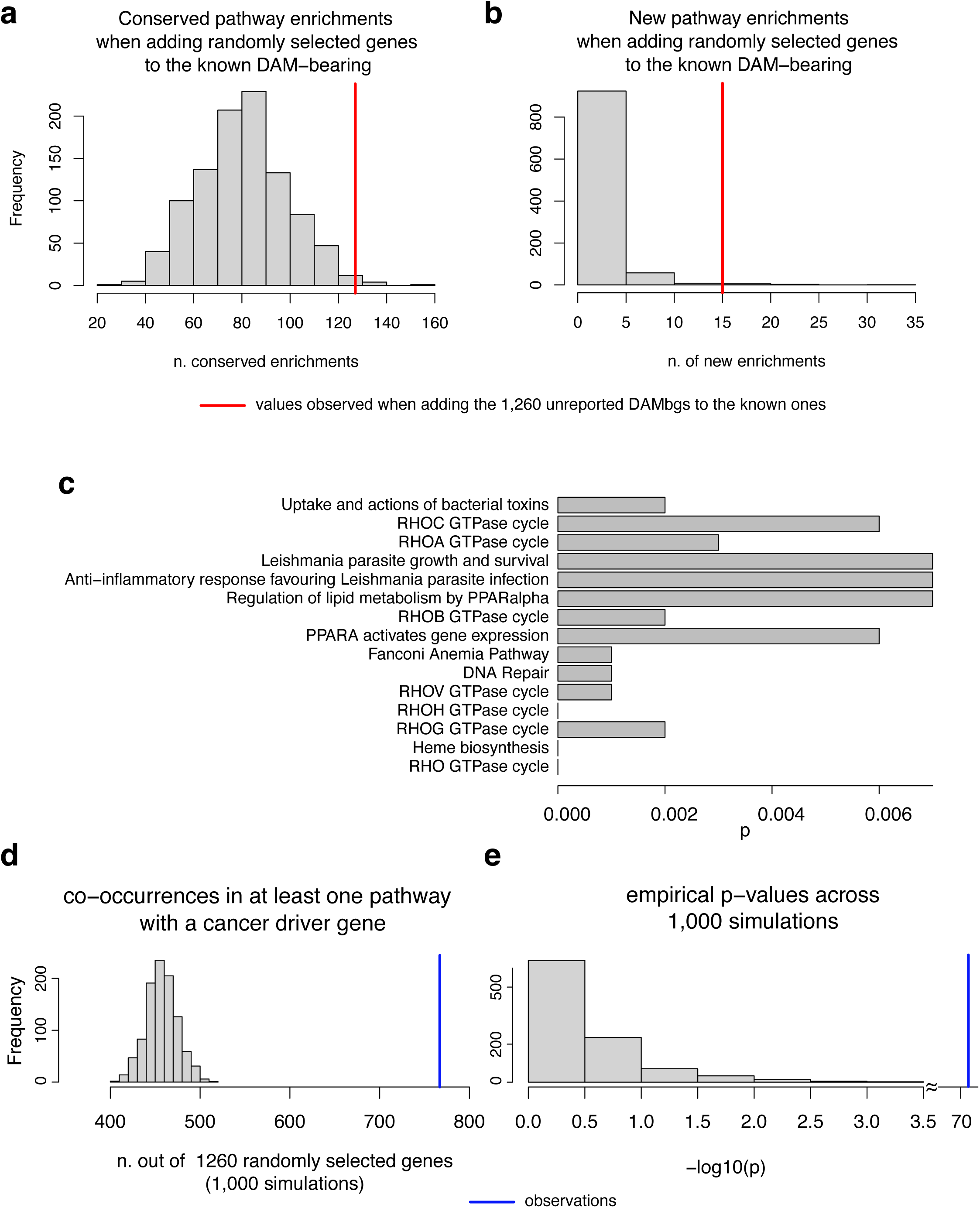
a-b. Number of pathway enrichments that are conserved (a) or gained (b) when adding to the known DAM-bearing genes (DAMbgs), 1,260 randomly selected genes that are not known to be cancer drivers, with the same ratio of genes that belong to at least one Reactome pathway as the unreported DAMbgs. The vertical red lines indicate observed values when adding the actual 1,260 unreported DAMbgs to the known ones. c. Frequency of observing the listed pathways as novel enrichments when adding to the known DAMbgs, 1,260 randomly selected genes that are not known to be cancer drivers and with the same ratio of genes that belong to at least one Reactome Pathway of the unreported DAMbgs, across 1,000 trials. d. Distribution of the number of genes that co-occur with a known cancer driver in at least one Reactome pathway (out of 1,260 randomly selected ones), across 1,000 samplings. e. Distribution of empirical Hypergeometric test p-values quantifying the probability of the numbers in d, across 1,000 simulations. Observed values are indicated by the vertical red lines.

**Extended Data Fig. 9.**
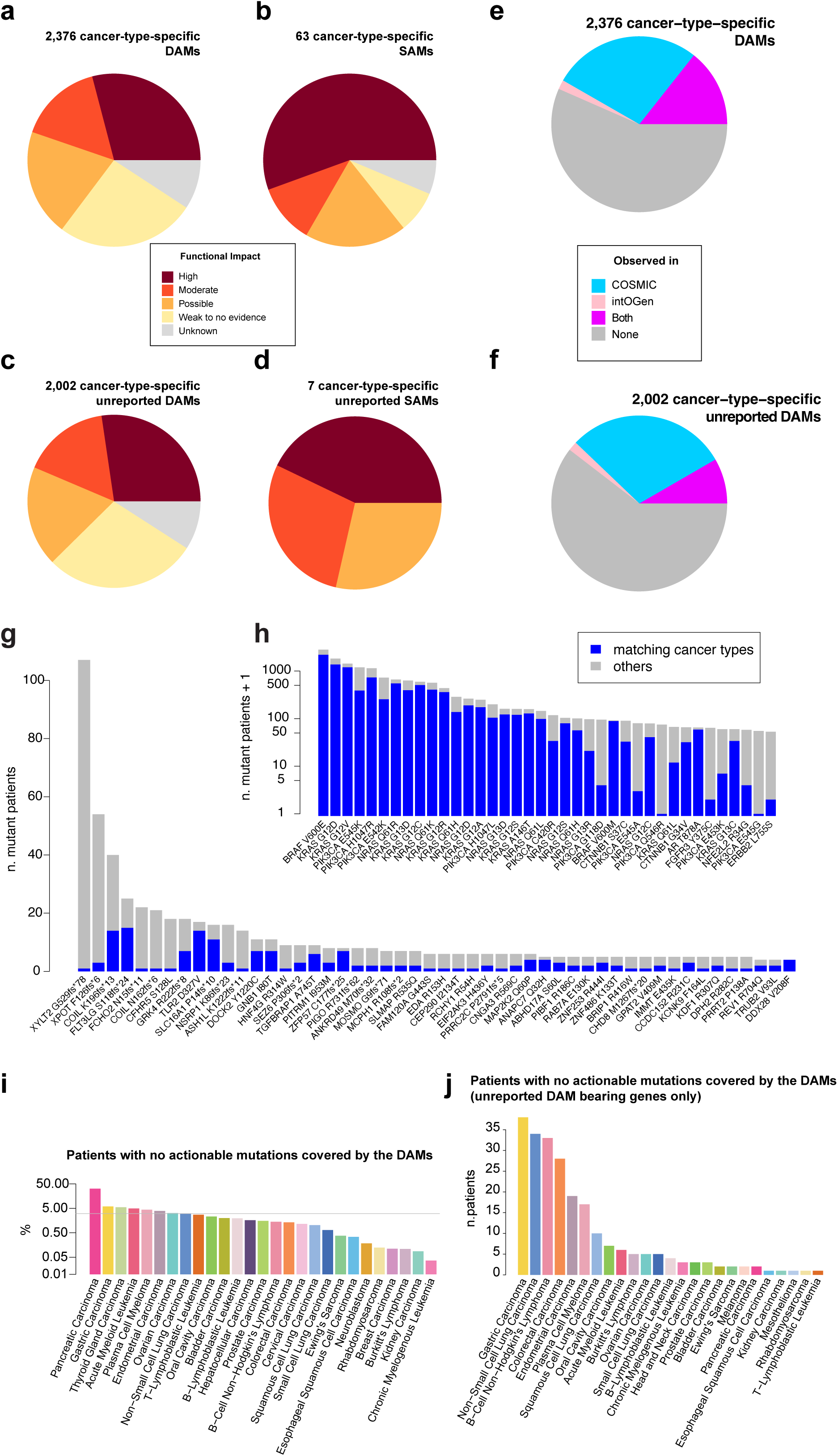
a-d. Collapsed predicted functional impact class for all Dependency-Associated Mutations (DAMs) and Drug-Sensitivity-Associated Mutations (SAMs, a-b) and for DAMs and SAMs involving genes not previously reported as cancer driver genes (c-d). e-f. Fractions of DAMs observed in at least one patient in publicly available cancer genomics resources, considering all of them (e) or only those involving genes not previously reported as cancer drivers (f). g-h. DAMs most frequently observed in cancer patients and involving established cancer driver genes (h) or genes not previously reported as cancer drivers (g). The blue bars indicate the fraction of patients with a cancer matching that in which the DAM has been identified. ii. Percentages of patients in COSMIC that harbour a DAM and no other variants that are currently clinically actionable, across analysed cancer-types. j. As in i. but considering the number of patients and only DAMs hosted by genes not previously reported as cancer drivers.

**Extended Data Fig. 10.**
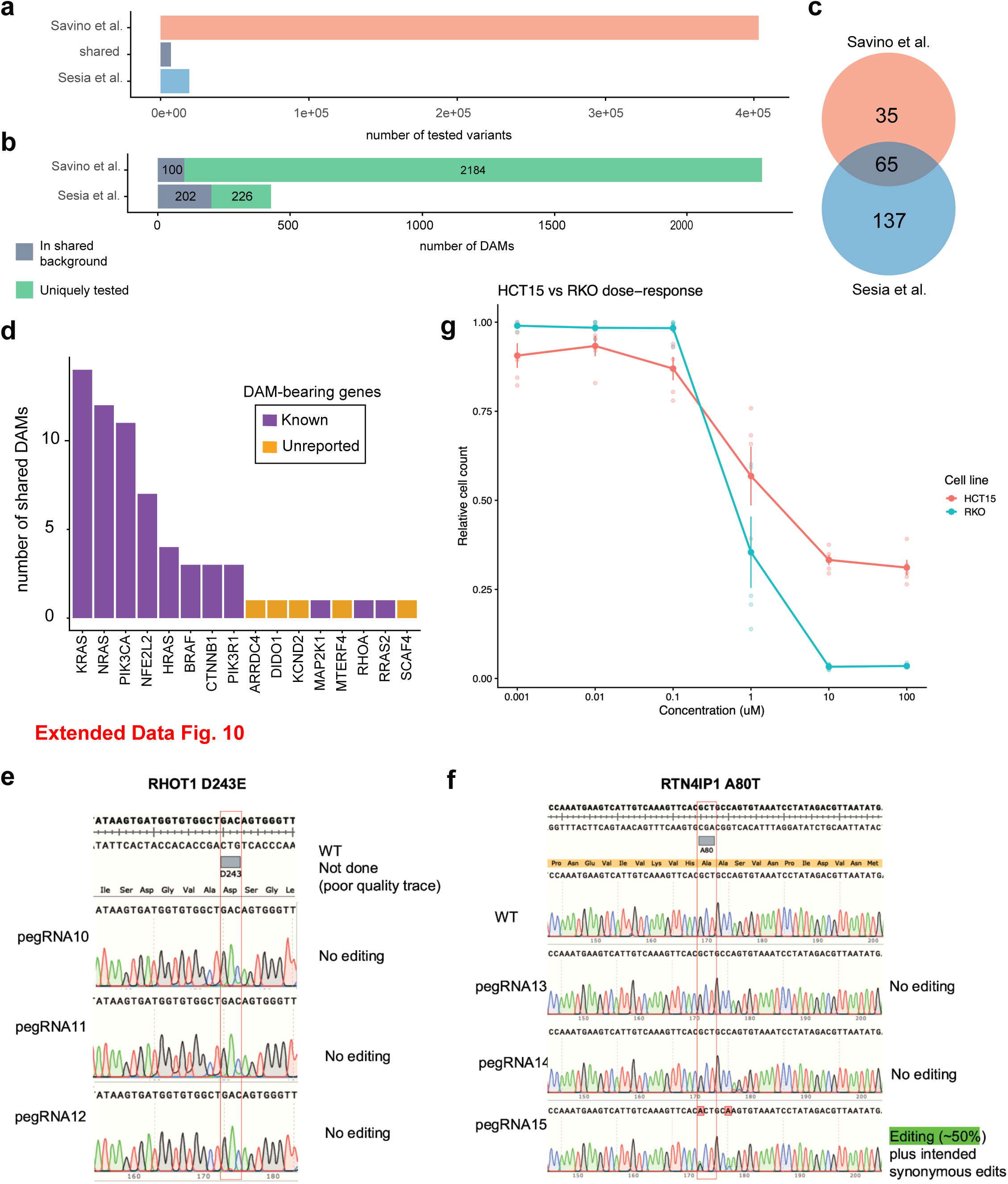
a-d. Independent analytical pipelines converge on shared Dependency-Associated Mutations. a. Numbers of variants tested by CRISPR-VUS (Savino et al.; 403,269 variants, including singleton events) and Sesia et al. (18,680 recurrent variants), together with their common 6,768 tested background. CRISPR-VUS tested variants against dependency on their host genes, whereas Sesia et al. also tested associations with dependencies on other genes. b. Numbers of DAMs identified by each study after collapsing the cancer-type-specific CRISPR-VUS results to unique variants. Purple segments denote DAMs within the common tested background (CRISPR-VUS, 100; Sesia et al., 202), whereas green segments denote DAMs involving variants tested only by the respective study (CRISPR-VUS, 2,184; Sesia et al., 226). c. Overlap between DAMs identified within the common tested background. The two studies shared 65 DAMs (substantially more than the 2.98 expected by chance, 21.8-fold enrichment; one-sided hypergeometric test, P = 4.89 × 10⁻⁷⁸), with 35 identified only by CRISPR-VUS and 137 only by Sesia et al. The background comprised all variants tested in both analyses. Cancer-type information was disregarded because Sesia et al. reported pan-cancer rather than tissue-specific associations. d. Distribution of the 65 shared DAMs across their host genes. Bar heights indicate the number of shared DAMs per gene; purple denotes established cancer-driver genes and orange denotes unreported DAM-bearing genes. Five shared DAMs mapped to five genes not previously classified as cancer drivers: ARRDC4, DIDO1, KCND2, MTERF4 and SCAF4. e-f. Sanger sequencing chromatograms of cells targeted to introduce the RHOT1 D243E (e) or RTN4IP1 A80T (f) variants. No detectable editing was observed with RHOT1 pegRNAs or RTN4IP1 pegRNAs13-14. RTN4IP1 pegRNA15 achieved approximately 50% editing and introduced the intended A80T variant together with the designed synonymous substitutions. g. Istaroxime dose-response profiles in ATP1B3 p.I189M-bearing and wild-type colorectal carcinoma cells. ATP1B3-wild-type HCT15 cells (salmon) and ATP1B3 p.I189M-bearing RKO cells (teal) were treated with the indicated concentrations of istaroxime for 72 h under 0.5% serum. Cell viability was measured using CellTiter-Glo and expressed as relative cell count normalised to the corresponding vehicle-treated control. Small translucent points represent experiment-level means, obtained by averaging four technical replicate wells at each concentration. Large filled points and connecting lines show the mean across five independent experiments; error bars indicate s.e.m. Lower relative cell counts indicate greater sensitivity to istaroxime. The fitted log₁₀IC50, area under the dose–response curve and Emax values derived from the individual experiments are summarised in Fig. 5b (n=5 independent experiments).

