## Supplementary Methods for "CRISPR-enhanced assessment of variants of unknown significance nominates oncology therapeutic targets and drug repositioning opportunities"

### *Input Data and the cancer cell lines (CCLs) analysis set: additional information*

Full provenance information for all input datasets mentioned below - including exact filenames, release versions, download URLs, download dates and content descriptions - is provided in **Supplementary Table 1**.

We assembled molecular profiles, functional-screening data and model annotations for cancer cell lines (CCLs) represented in the Cell Model Passports (CMP)[1]. We downloaded the CMP model catalogue (`model_list_20241120.csv`) on 29 January 2025. The catalogue contained 2,161 models, of which 1,430 had available somatic-variant calls in `mutations_all_20241212.zip`, downloaded on 21 February 2025. We obtained gene-expression profiles, expressed as fragments per kilobase of transcript per million mapped reads (FPKM), from the CMP RNA-sequencing dataset `rnaseq_all_20220624.zip`. We downloaded `gene_identifiers_20241212.csv` on 21 February 2025 and used it to harmonise gene identifiers across datasets.

We obtained CRISPR gene-effect scores from the DepMap 24Q4 release (`CRISPRGeneEffect.csv`), downloaded on 29 January 2025, harmonised (across Broad and Sanger CRISPR datasets) and scaled as we previously described[2,3] and applied the quality-control procedure that we previously designed[4]. We excluded models absent from the CMP model catalogue.

As reference sets of core-fitness and common-essential genes, we used our previously published lists from Vinceti et al.[5], computed with the Adaptive Daisy Model (AdAM) and the FiPer-AUC method, respectively.

After dataset matching, quality control and cancer type selection, we considered 36 different cancer types, for which at least five CCLs were available. We also excluded CCLs assigned to the non-specific categories “Other Solid Carcinomas”, “Other Solid Cancers”, “Other Sarcomas” and “Other Blood Cancers”, as well as models annotated as “Non-Cancerous”.

Following this model matching and quality control, the analysis included 977 CCLs across 36 cancer types (the CCL analysis set, with a median number of CCLs per cancer type = 24, min = 6 for Burkitt's Lymphoma, max = 69 for Non-Small Cell Lung Carcinoma, **Supplementary Table 1** and **Extended Data Fig. 1a**).

We obtained drug-response measurements from the GDSC[6], release 8.5 datasets. We downloaded the fitted dose–response files on 3 October 2024 and used the reported half-maximal inhibitory concentrations (IC50s) for the pharmacological analyses.

We obtained cancer-driver annotations from the IntOGen[7] driver compendium released on 20 September 2024, downloaded on 2 October 2024, and from additional benchmark cancer-driver datasets compiled in **Supplementary Table 1**. We obtained target-tractability annotations from the OpenTargets[8] Platform tractability dataset downloaded on 29 August 2025. We used these annotations only after DAM discovery to support biological interpretation and translational prioritisation.

For patient-level analyses, we obtained somatic-variant and sample-annotation data from COSMIC[9]. We used the GRCh38 Genome Screens Mutant, Sample and Classification datasets from COSMIC v101 and COSMIC v104, downloaded on 13 July 2026. We obtained clinical-actionability annotations from the CIViC[10] Clinical Evidence Summaries released on 1 May 2025 and downloaded on 16 May 2025.

We used CMP model identifiers as the primary identifiers for CCLs and mapped DepMap and GDSC identifiers to CMP models using the cross-references and model synonyms provided in `model_list_20241120.csv` (downloaded from the CMP). We harmonised genes to approved HGNC symbols using `gene_identifiers_20241212.csv` (downloaded from the CMP), which provides mappings among HGNC, Ensembl, Entrez, RefSeq, UniProt and COSMIC identifiers. We matched variants across datasets using harmonised gene symbols and protein-level variant annotations. We harmonised cancer type labels across CMP, IntOGen, COSMIC and CIViC using manually curated mapping tables (**Supplementary Table 1**).

We obtained gene-level loss-of-heterozygosity (LOH) calls from the DepMap Public 24Q4 release (OmicsLoH.csv), downloaded on 17 July 2026. This dataset provides a binary gene-by-cell-line matrix indicating the presence or absence of LOH for each gene in each model. We harmonised DepMap model identifiers with CMP identifiers as described above. We used these data only for the post-discovery characterisation of frameshift DAMs and not for DAM identification or threshold selection.

***Selection of somatic variants for dependency testing: additional information***

The somatic-variant catalogue obtained from the CMP had already undergone germline-variant removal and quality control. Within each cancer type, we grouped all occurrences of the same variant using the harmonised host-gene symbol and protein-level variant annotation. We treated the occurrence of a variant in a particular cancer type as an individual testable variant/cancer type combination. Consequently, a variant observed in more than one cancer type contributed one testable combination for each cancer type, whereas multiple occurrences of the same variant among CCLs belonging to the same cancer type contributed multiple variant instances to a single combination.

We first excluded genes previously classified as core-fitness genes or common essentials[5]. Because these genes are required for the viability of most proliferating cells, their knockout produces broadly deleterious effects largely independently of the presence of a particular somatic allele. Their inclusion would therefore have provided limited information about allele-specific genetic dependencies and could have generated associations driven by constitutive gene essentiality.

We then restricted the analysis to protein-altering coding variants that could plausibly generate allele-specific functional consequences. We retained missense substitutions, in-frame insertions or deletions and frameshift variants. Missense and in-frame variants can alter protein activity, interactions, localisation or regulation without necessarily eliminating the encoded protein and therefore represent plausible sources of gain-of-function, neomorphic, dominant-negative or separation-of-function phenotypes.
Although frameshift mutations frequently cause loss of function, we retained them because some frameshifts can produce stable truncated proteins, recurrent alternative protein products, partial loss-of-function states or other allele-specific phenotypes associated with selective genetic dependencies[11,12]. We subsequently evaluated the plausibility of frameshift DAMs using their variant allele fractions and loss-of-heterozygosity patterns, rather than excluding the entire mutation class a priori.

We excluded variants annotated as synonymous or silent, nonsense or stop-gain, start-loss, stop-loss, or essential splice-site variants. Synonymous variants were excluded because they do not ordinarily alter the encoded protein sequence, whereas the remaining classes predominantly generate complete gene inactivation, unstable transcripts or heterogeneous transcript-level consequences. Such effects are less readily attributable to a specific altered protein allele and are therefore less suited to the allele-specific dependency framework used here. Variants outside the retained missense, in-frame insertion/deletion and frameshift classes were not tested.

We additionally excluded, within each cancer type-specific analysis, genes harbouring more than ten distinct variants across the corresponding CCLs. A broad and heterogeneous mutational spectrum is more characteristic of genes undergoing non-selective mutation accumulation or recurrent loss of function than of genes affected by a restricted set of allele-specific activating alterations. This filter also reduced the contribution of very long genes, such as TTN, which accumulate numerous passenger mutations because of their coding length. The filter was applied to the number of distinct qualifying variants in a gene, rather than to the total number of variant-bearing CCLs.

We did not require variants to recur across CCLs. Singleton variants, i.e. observed in only one CCL within a cancer type, were retained and evaluated alongside recurrent variants. This was essential because singleton and near-singleton events constituted the majority of the eligible variant catalogue and represented a principal target of the CRISPR-VUS framework.

Cancer-driver status, predicted functional impact, occurrence or recurrence in patient tumours, target tractability and pharmacological annotations were not used to select variants for dependency testing. Similarly, we did not restrict the discovery analysis to genes expressed above a predefined threshold. These features were incorporated only after DAM identification as orthogonal annotations for biological interpretation, validation and translational prioritisation. Retaining variants in non-expressed genes also enabled their use as an empirical decoy set for the independent estimation of the global false discovery rate.

After applying the model and gene and variant-class filters, the discovery space comprised 532,331 testable variant/cancer type combinations (**Figure 1d**), corresponding to 403,269 distinct somatic variants in 16,842 genes. Because some variants occurred in more than one CCL within the same cancer type, these combinations represented 585,344 individual variant/CCL instances. This complete filtered set constituted the input to the subsequent rankRatio-based dependency analysis and is provided in **Supplementary Dataset 1**.

#### *Analysis of DAMs in established drivers outside their recognised cancer contexts, canonical-hotspot proportions: additional information*

We classified a DAM-bearing gene as an established cancer driver when it appeared in the IntOGen[13] cancer-driver compendium. We determined the recognised cancer contexts of each driver from the cancer-type-specific IntOGen annotations. Cancer-type labels were harmonised between IntOGen and the Cell Model Passports using the manually curated mappings provided in **Supplementary Table 1**.

We evaluated each DAM–cancer-type association separately. We classified an association as off-context when the DAM occurred in an established cancer-driver gene but IntOGen did not identify that gene as a driver in the corresponding cancer type or an equivalent mapped tumour category. Thus, "off-context" indicates the absence of a previously reported driver association for that gene/cancer-type combination and does not imply that the gene or variant lacks functional relevance in that context.

To quantify the occurrence of off-context DAMs in patient tumours, we matched their protein-level annotations to COSMIC variants as described above. For each gene/cancer-type combination, we calculated the percentage of analysed patients in the corresponding COSMIC tumour cohort carrying at least one matching off-context DAM. Patients carrying multiple qualifying DAMs in the same gene were counted once for that gene/cancer-type combination.

We next assessed whether off-context DAMs nevertheless corresponded to recognised oncogenic hotspot alterations. The 187 cancer-type-specific DAM associations involving established drivers outside their recognised contexts were cross-referenced against the IntOGen driver-mutation catalogue. Variant annotations were harmonised at the protein level before matching. We classified a DAM as a canonical hotspot when its gene and amino-acid alteration exactly matched an IntOGen driver mutation. Variants without an exact match were classified as non-canonical. The proportions of canonical and non-canonical alterations were calculated across the 187 off-context DAM associations.

#### *Cell-line misclassification controls: additional information*

Because apparent off-context dependencies could arise from incorrect cell-line annotations, cross-contamination or ambiguous histopathological origins, we performed a multi-omic cell-line similarity analysis. We considered every CCL supporting an off-context DAM and compared its molecular profile with those of the other models across four independently derived data modalities: gene expression, protein abundance, CRISPR-based gene dependency and drug response. The target CCL itself was excluded from its reference set.

Within each modality, we calculated Pearson correlations between the profile of the CCL under consideration and the profiles of all other models with sufficient overlapping measurements. We then summarised its similarity to models belonging to its annotated cancer type and to each alternative cancer type in which the corresponding DAM-bearing gene was recognised as a driver. When a gene was an established driver in more than one alternative cancer type, each alternative context was evaluated separately.

To test whether a CCL showed greater similarity to an alternative lineage than expected by chance, we randomly permuted the cancer-type labels of the reference models 1,000 times while retaining their molecular profiles unchanged. After each permutation, we recalculated the cancer-type similarity scores. We derived a one-sided empirical P value from the proportion of permutations in which the similarity to the alternative cancer type equalled or exceeded the observed similarity. We examined the results across all available modalities to identify models showing consistent evidence of molecular resemblance to an alternative lineage rather than relying on a single data type.

This analysis served as a diagnostic control for possible cell-line misclassification. It was performed after DAM discovery and did not alter the cancer-type annotations, DAM calls or thresholds used by CRISPR-VUS.
