## Supplementary Results for "CRISPR-enhanced assessment of variants of unknown significance nominates oncology therapeutic targets and drug repositioning opportunities"

### Additional quantitative evidence supporting the biological plausibility of DAMs

Frameshift Dependency-Associated Mutations (DAMs) exhibited genomic characteristics distinct from other mutation classes. Their variant allele fractions were significantly lower than those of missense and in-frame DAMs (two-sided Wilcoxon rank-sum test, $P = 2.05 \times 10^{-9}$; Cliff's $\delta = 0.26$; **Extended Data Fig. 2e**). Frameshift DAMs also showed a lower prevalence of loss of heterozygosity (LOH) than background tested frameshift variants, whereas no comparable difference was apparent for missense and in-frame variants. However, the variant-class-by-LOH interaction did not reach statistical significance ($P = 0.10$; **Extended Data Fig. 2f**). These observations are therefore compatible with, but do not establish, heterozygous contexts in which partial disruption of one allele increases reliance on the remaining functional activity of the host gene[1].

DAM-bearing genes were frequently highly expressed within the CCLs carrying the corresponding variants. Among DAM instances, 92% involved genes expressed above the 50th percentile within the host CCL, 62% above the 80th percentile and 16% above the 95th percentile (**Fig. 2b**). Consistently, logistic regression showed that basal expression percentile strongly predicted DAM classification: every 10-percentile increase in expression was associated with a 34% increase in the odds that a tested variant instance was identified as a DAM (odds ratio = 1.34, 95% CI 1.312–1.37, $P < 9.95 \times 10^{-152}$; **Extended Data Fig. 2g**). Thus, DAM enrichment was not restricted to genes marginally exceeding the FPKM threshold but increased progressively across the cellular expression distribution.

The decoy-based false-discovery analysis also quantified the contribution of the HIT-local statistical filters. At the operational rankRatio threshold of 1.71, the estimated global FDR was 23% before application of the permutation and hypergeometric filters and 22% afterwards. The filters removed 6% of predicted DAM instances but 11% of decoy DAM instances, indicating a modest but measurable improvement in specificity (**Extended Data Fig. 2ac**).

### Cancer-type variation and robustness against established driver catalogues

The number of identified DAMs varied substantially among cancer types. The median was 57.5 DAMs per cancer-type-specific analysis, ranging from 7 in T-cell non-Hodgkin lymphoma to 245 in colorectal carcinoma. The number of DAM-bearing genes ranged from 7 to 167, with a median of 49.5 (**Supplementary Tables 2-3** and **Extended Data Fig. 1cd**). Although absolute DAM numbers increased with the number of available CCLs and tested variants, the normalised signal-to-noise analysis showed that this did not correspond to an increased proportion of positive predictions in more highly mutated cancers (**Fig. 2fg**, **Supplementary Table 1**, **Extended Data Fig. 3b-d**).

The overlap between DAM-bearing genes and established cancer-driver catalogues was observed separately across missense, frameshift and in-frame variants, indicating that recovery of recognised cancer dependencies was not exclusively driven by the predominant missense class (**Extended Data Fig. 2d**). Among the 123 IntOGen driver genes identified as DAM-bearing, 60 (49%) were annotated as harbouring both activating and loss-of-function driver alterations in different cancer contexts (hypergeometric test, $P = 8.8 \times 10^{-74}$; **Extended Data Fig. 4e**). In the cancer-type-specific analyses, enrichment was more frequently observed for drivers with recognised gain-of-function roles than for genes acting predominantly through loss of function (**Extended Data Fig. 4f**). Conversely, 220 of the 290 analysable IntOGen drivers not recovered as DAM-bearing genes (76%) were classified as tumour suppressors or had ambiguous functional roles (**Extended Data Fig. 4d**). These results are consistent with preferential detection of variants that generate or reinforce dependency on their host genes.

#### Recurrently identified unreported DAM-bearing genes

Among the 332 DAM-bearing genes identified in multiple cancer types, several unreported genes - including ALG10B, DIDO1, CHTF18, TRMT6 and TRUB2 - had previously been implicated in cancer-associated cellular processes (**Extended Data Fig. 5a**). Their repeated identification across independent cancer-type-specific analyses supports their prioritisation as candidate context-dependent dependencies.

RIF1 provided a particularly notable example. RIF1 DAMs were identified across six cancer-type-specific analyses and distributed across two broad functional regions. Variants p.V133A, p.D495Y, p.S566L, p.Q926K/L, p.T986A and p.A1142T occurred within the N-terminal and central HEAT-repeat-containing regions involved in chromatin association and replication timing. A second group, comprising p.E1620K, p.A2031G, p.D2108G and p.C2223F, mapped to the C-terminal region implicated in the 53BP1-RIF1 DNA-repair axis. These alterations were missense substitutions, several involving marked physicochemical changes that could perturb specific protein interactions without eliminating the protein. The p.E1620K variant contributed to more than one DAM hit, while p.E1620K and p.D495Y were also observed in patients with tumour histologies matching the CCL contexts in which they were identified (**Supplementary Table 5**). Although these observations do not establish a shared molecular mechanism, they provide additional support for investigating RIF1 as a context-specific dependency associated with replication-stress tolerance and DNA-repair pathway choice.

A smaller number of recurrently detected established DAM-bearing genes, including TSC2, ARID1A and PBRM1, have canonical tumour-suppressor roles. Most of the DAMs identified in these genes were missense rather than truncating variants. One possible interpretation is that these alleles preserve residual or altered protein activity upon which the cells remain dependent. However, this hypothesis requires direct experimental investigation, and these associations should not be interpreted as evidence that the variants activate the corresponding tumour suppressors.

#### DAMs in established drivers outside their recognised cancer contexts

187 cancer-type-specific DAMs involved established cancer-driver genes outside the tumour contexts in which IntOGen currently recognises them as drivers (**Extended Data Fig. 5b**). Of these, 23 (12.3%) corresponded to canonical driver hotspots, whereas 164 (87.7%) did not (**Supplementary Table 3**). The canonical-hotspot group involved NRAS, BRAF, PIK3CA, HRAS, ERBB2, MAP2K1, RRAS2, NFE2L2, CREBBP, EP300, PIK3CB and SMARCA4. Clear examples included the NRAS codon-61 substitutions p.Q61K, p.Q61L and p.Q61R, identified as DAMs in B-cell non-Hodgkin lymphoma and bladder carcinoma.

These off-context associations may reflect an incompletely characterised oncogenic role across tumour lineages or an uneven representation of molecular subtypes in available CCL collections. Alternatively, apparent off-context associations could arise from incorrect lineage annotation, cross-contamination or ambiguous histopathological origin. To examine the latter possibility, we compared the implicated CCLs with other models using transcriptomic, proteomic, CRISPR-dependency and pharmacological profiles. This analysis provided little evidence of systematic reassignment of these models to lineages in which the corresponding genes were established drivers (**Extended Data Fig. 5c-h**).

DAMs were also not concentrated in a small subset of potentially problematic models. They occurred across 596 CCLs, corresponding to 61% of the analysed collection, and most DAM-bearing CCLs contained no more than three DAMs (**Extended Data Fig. 6a-c**). This broad distribution argues against widespread misclassification or a small number of highly mutated CCLs accounting for the observed associations.

### Additional pathway-level observations

Unreported DAM-bearing genes did not produce statistically significant pathway enrichments when analysed in isolation (**Supplementary Table 3**). Their contribution instead became evident when they were added to the established DAM-bearing genes, suggesting that they populate additional nodes within recognised cancer programmes rather than forming a separate group of highly recurrently altered pathways.

Variant-level examples illustrate this incremental contribution. In ERK signalling, MAP2K2 p.Q60P in plasma cell myeloma, MAPK14 p.E253A and p.L289F in gastric carcinoma, and RAPGEF1 p.R1051W in T-lymphoblastic leukaemia complemented established DAM-bearing genes such as BRAF, KRAS, MAP2K1 and NRAS. Within the PI3K/AKT cascade, CHUK p.E496D in B-cell non-Hodgkin lymphoma and IRS2 p.E1242V in colorectal carcinoma complemented PIK3CA, PIK3CB, EGFR and ERBB2. Similarly, BRAP DAMs in B-lymphoblastic leukaemia and prostate cancer, CSK p.Y403C in head and neck carcinoma, IQGAP1 variants in NSCLC and gastric carcinoma, and SRC variants in colorectal and pancreatic carcinoma extended networks associated with mutant RAS signalling (**Fig. 3e-g**).

Several of the pathways that became significant only after adding the unreported DAM-bearing genes have established roles in tumour biology. RHO-family GTPase cycles regulate cytoskeletal organisation, migration and invasion; DNA-repair and Fanconi Anaemia pathways generate therapeutically relevant genome-maintenance dependencies; and PPARα-regulated pathways implicate lipid metabolism and inflammatory signalling in cellular adaptation. The emergence of these enrichments only after inclusion of unreported DAM-bearing genes suggests that rarely mutated components can collectively extend the detectable architecture of established cancer pathways.

### Additional patient-cohort observations

Of the DAMs observed in patient tumours, 341 were represented in both COSMIC and IntOGen. Among DAMs involving unreported DAM-bearing genes, 167 were found in both resources (**Extended Data  Fig. 9e-f** and **Supplementary Table 5**). Furthermore, among the 675 DAMs that were both predicted to affect protein function and observed in patients, 468 involved unreported DAM-bearing genes. Of the 352 DAMs observed in histology-matched patient tumours, 197 involved unreported genes (**Fig. 1d** and **Supplementary Table 2**).

Examples of recurrent patient-observed DAMs in unreported genes included *FLT3LG p.S118fs**24 in colorectal carcinoma, COIL p.K196fs*13 in gastric carcinoma, TLR2 p.D327V in B-cell non-Hodgkin lymphoma and XYLT2 p.G529fs**78* in small-cell lung carcinoma (**Fig. 4a** and **Supplementary Tables 5-6**). These findings show that variants appearing as singleton or near-singleton events in CCL panels can nevertheless recur in patients, including within the histological context in which their dependency association was detected.

Among the 7,190 COSMIC patients carrying at least one DAM, 2,796 lacked any co-occurring clinically actionable variant annotated in CIViC (**Fig. 4b**). When restricting the analysis to DAMs in unreported genes, gastric and endometrial carcinomas contained identifiable patient subgroups lacking established actionable alterations. These included 38 gastric carcinoma patients and 19 endometrial carcinoma patients. Recurrent examples included COIL p.K196fs**13 in gastric carcinoma and PIGO p.R731fs**62, EIF2AK3 p.R114I and WRAP53 p.R194Q in endometrial carcinoma (**Fig. 4b** and **Extended Data Fig. 9ij**).

Patient data also supported some off-context associations involving established drivers. NRAS DAMs were observed in additional tumour contexts including B-cell non-Hodgkin lymphoma, bladder carcinoma and neuroblastoma. JAK3 DAMs occurred in a subset of acute myeloid leukaemia patients, while BRAF DAMs occurred in Ewing sarcoma patients (**Fig. 4c**). These findings nominate possible extensions of established driver dependencies to additional tumour contexts but do not independently demonstrate clinical actionability.

### Complete pharmacological-screening results and illustrative repurposing hypotheses

The pharmacological analysis evaluated 4,017 drug–target/cancer-type combinations involving 103 compounds and 31 target genes. Among the 1,899 cancer-type-specific dependency hits, 74 involved a druggable DAM-bearing gene targeted by at least one compound with matched GDSC response data. Twenty-nine of these 74 hits (39%) met the SAM criteria, collectively encompassing 63 individual Sensitivity-Associated Mutations and 33 compounds (**Fig. 1d** and **Supplementary Table 2**). These denominators demonstrate the extent of the pharmacological search space from which the SAMs were identified.

Several actionable DAMs in unreported genes suggested hypotheses for pharmacological investigation. Examples included TLR2 p.D327V in B-cell non-Hodgkin's lymphoma; NDUFS1 p.A594S and p.V624G, NCSTN and AVPR1B variants in colorectal carcinoma; KCNK9 p.R142P and p.F164L in lung cancer; and SV2A p.S281C in squamous carcinoma. These associations connect genetic dependencies to proteins already targeted by approved, clinical or investigational compounds. They should nevertheless be interpreted as prioritisation hypotheses rather than evidence of therapeutic efficacy, particularly when no matched public drug-response data were available.

MAP2K2 p.Q60P in plasma cell myeloma represented a more extensively supported example. In addition to its association with MAP2K2 dependency, the variant was consistently associated with greater sensitivity to PD0325901, refametinib and trametinib. It was also predicted to affect protein function and was observed in both COSMIC and IntOGen, including in the corresponding tumour context. The variant maps near the N-lobe of the MAP2K2 kinase domain and has previously been associated with increased ERK phosphorylation. Together, these observations support a model in which MAP2K2 p.Q60P marks a targetable MAPK-dependent state in plasma cell myeloma, while requiring prospective validation before clinical interpretation.
